# Targeted, high-resolution sensing of volatile organic compounds by covalent nanopore detection

**DOI:** 10.1101/2024.10.15.618413

**Authors:** Lauren E McGivern, Zhong Hui Lim, Yizhi Yuan, Zonghua Bo, Hagan Bayley, Yujia Qing

## Abstract

Volatile organic compounds are choice analytes in a variety of contexts. For example, humans release over 4000 volatile organic compounds, many of which are diagnostic of life-threatening medical conditions. A combination of a large number of potential analytes requires the application of costly, cumbersome technology. Here, we show that covalent nanopore sensing can be used for the targeted detection of a reduced set of analytes in a mixture: in this case aldehydes, which constitute ∼5% of human volatiles. Further, nanopore engineering permits high-resolution detection, which allows closely related aldehydes including isomers to be distinguished. Differential sensing of other chemical classes, such as alcohols, is demonstrated by leveraging their enzymatic conversion to aldehydes. Our approach is compatible with the use of cheap, portable, user-friendly diagnostic devices applicable to a wide variety of objectives, including pollutant monitoring, food and beverage testing and the quality control of pharmaceuticals, as well as disease diagnostics.

## Introduction

Aldehydes are ubiquitously produced through chemical processes across various biological, environmental, and industrial contexts. These include lipid peroxidation in the human body,^1^ incomplete combustion during wood burning,^2^ and Strecker degradation during beer storage.^3^ Single-molecule detection of diverse aldehydes against background chemicals therefore offers wide applications.

For example, small-molecule aldehydes are found in human breath or bodily fluids, alongside a spectrum of volatile organic compounds (VOCs) generated through metabolic processes^1^ or by the gut microbiome.^4^ Metabolic alterations in specific conditions, such as cancers,^5^ gastrointestinal diseases,^6^ respiratory disorders,^7^ and viral infections,^8^ lead to distinct VOC profiles, which offer a non-invasive window into the body’s internal biochemical processes. The current gold standard for detection of small molecules is liquid or gas chromatography-mass spectrometry (LC/GC-MS),^9^ which generates a near-complete profile of all collected VOCs, but requires centralized labs that use expensive equipment and sophisticated analysis packages.

Here, we focus on the development of a single-molecule, real-time detection technology for rapid ratiometric profiling of aldehydes. The targeted detection of aldehydes, comprising about 170 species among over 4000 VOCs of human origin,^10^ presents an appealing approach to reduce the complexity of small-molecule fingerprints, while retaining significant diagnostic value. Aldehyde mixtures in breath or bodily fluids commonly contain linear or branched, unsaturated or saturated carbon chains, containing from 1 to 17 carbon atoms, as well as aromatic species such as benzaldehyde and its derivatives.^10^ A ratiometric fingerprint of aldehydes holds promise for disease detection and health monitoring, provided that tests can be made rapid, simple and inexpensive. For example, the ethanal:pentanal:heptanal ratio in breath shifts from 112:1:1.5 in healthy individuals to 24:1.7:1 in patients with lung cancer^11^, and similarly, the hexanal to heptanal ratio in urine changes from 1.3:1 to 2.5:1.^12^ For COVID infection, the relative abundance of octanal and benzaldehyde is a hallmark of recent infection.^8^

We have established single-molecule covalent sensing mediated by protein nanopores to detect analytes based on chemical reactivity.^13^ The concept was first demonstrated with engineered α-hemolysin (αHL) pores^14-17^ and later validated by others using alternative nanopores^18-20^. Analyte molecules form covalent bonds reversibly or irreversibly with a sensing group on the internal surface of a pore, thereby generating characteristic changes in the ionic current flowing through the pore under a transmembrane potential. When examining mixed analytes, the current signatures (e.g., the extent of current blockade and the noise during blockades) identify analytes, while the frequency of reversible events reveals the concentration of individual analytes.^21^

One general challenge is to identify sensing chemistry with suitable kinetics to ensure rapid and quantitative detection of analytes; the ideal lifetimes of covalent analyte-pore adducts and the intervals between sensing events are tens to hundreds of milliseconds, which can be reliably measured by electrical recording. This requirement has so far limited the practical application of covalent sensing. Another challenge is the rational engineering of nanopore sensors to resolve structures with minor variations, such as chain isomers that differ by the position of a methyl group. To date, this has largely relied on trial and error. A more systematic investigation is essential to guide the future development of covalent sensors.

Here, we apply such a systematic approach, to exploit hemithioacetal chemistry for the covalent sensing of aldehydes within a thiol-containing αHL nanopore, achieving rapid, high-resolution analysis by the formation of short-lived adducts that are identified by a machine learning algorithm. The diastereomeric adducts produce current signatures with differences that are accentuated by rational engineering of nanopores, affording information that discriminates between closely similar molecular isomers. We extend the scope of our approach to alcohols by selectively converting them to aldehydes. Hence, we have developed a versatile means for the targeted detection of a subset of analytes present within a complex mixture of molecules.

## Results and discussion

### Covalent detection of aldehydes through reversible hemithioacetal formation

Stochastic sensing of aldehyde analytes was achieved using reversible thiol-aldehyde chemistry. To eliminate background reactions, notably imine formation and metal chelation, four mutations were introduced in wild-type αHL (i.e., AG = K8A-M113G-K131G-K147G) (Supplementary Fig. 1).^22^ A single-cysteine mutation was then introduced at position 115 (i.e., AG-T115C). Heteroheptameric nanopores containing one cysteine-bearing subunit, (AG)_6_(AG-T115C), were prepared and used for covalent detection (Fig. 1a, see Methods). (AG)_6_(AG-T115C) carried a single-channel current (I_P_) of −129 ± 3 pA at −50 mV (N = 80 pores) (recording buffer: 2 M KCl, 200 mM PIPES and 20 μM EDTA at pH 6.8).

**Fig. 1:**
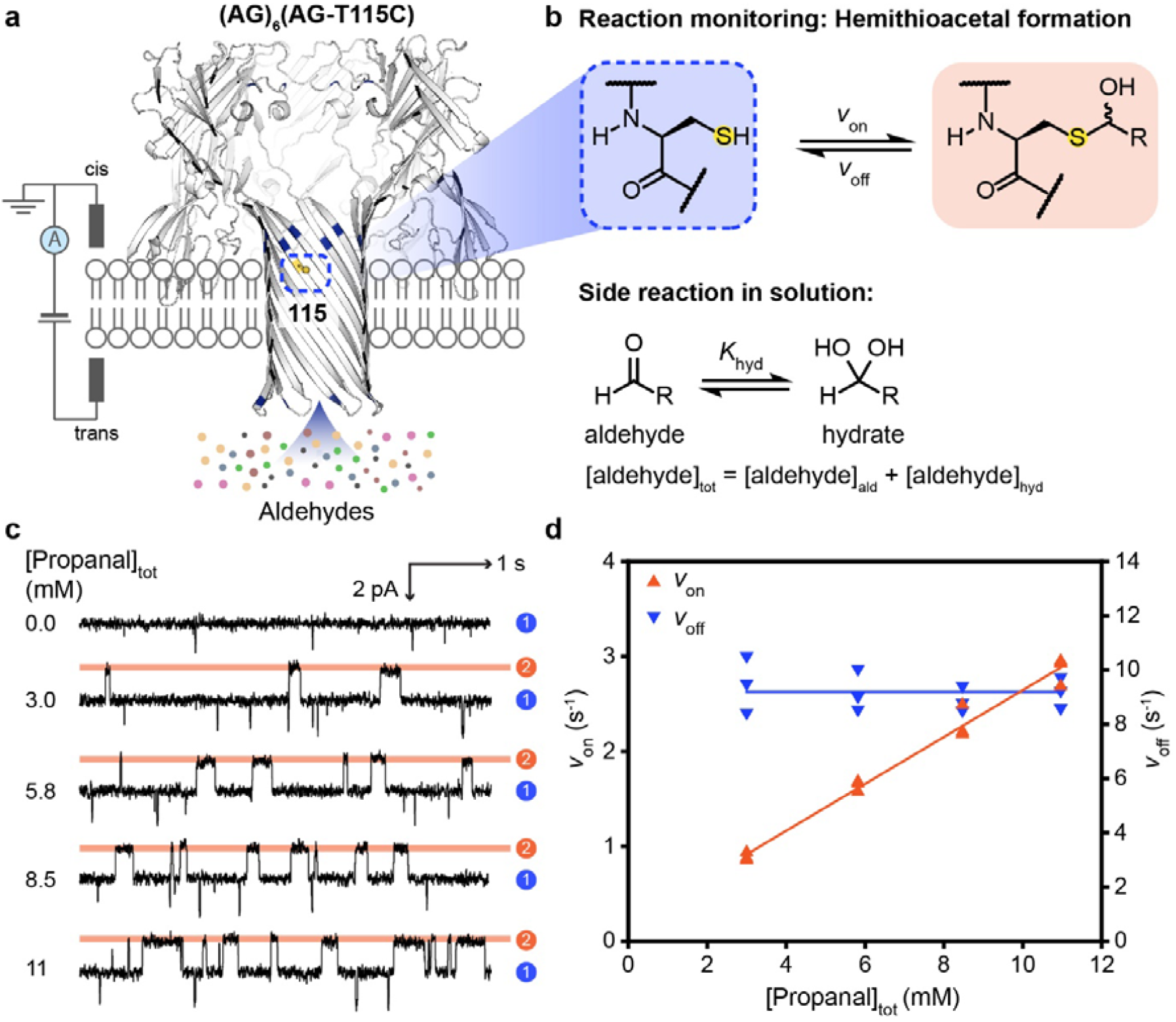
Single-molecule covalent sensing of aldehydes. **a**, The engineered α-hemolysin (αHL) nanopore, (AG)_6_(AG-T115C), bears a single cysteine at position 115 on one of the seven subunits (boxed). **b**, An aldehyde molecule was detected upon reaction with the cysteine thiol to form a hemithioacetal adduct. Aldehydes are hydrated to varying extents in aqueous solutions. Concentrations of free aldehyde were calculated by using the hydration constant K_hyd_ determined by ^1^H NMR. **c**, Single-channel recordings at −50 mV (trans) with 0.0, 3.0, 5.8, 8.5 or 11 mM propanal (trans) in 2 M KCl, 200 mM PIPES and 20 μM EDTA at pH 6.8. Signals were low-pass filtered at 10 kHz and sampled at 50 □ kHz. Traces were further filtered at 100 Hz for display. Current levels correspond to the hemithioacetal adducts formed (level 2, orange) and the unoccupied nanopore (level 1, blue). **d**, Rates of adduct formation (*v*_on_) and dissociation (*v*_off_), recorded with individual pores, plotted against total propanal concentration showing bimolecular kinetics for the forward association step (level 1 to 2), and unimolecular kinetics for the reverse dissociation step (level 2 to 1).

The introduction of an aldehyde in the trans compartment (Fig. 1a) produced reversible current blockades, which were attributed to the formation of hemithioacetal adducts (Fig. 1b,c and Supplementary Figs. 2-10). Residual currents (I_res_) after hemithioacetal formation are given as percentages of the open pore current (I_res%_ = I_res_/I_P_ × 100 %) (Supplementary Table 1). For example, for propanal, I_res%_ = 98.7 ± 0.1 % (N = 3 pores, >300 events). Aldehydes exist in both hydrated and non-hydrated forms in aqueous solution.^23^ The rates of hemithioacetal formation (*v*_on_) were expressed in terms of the total aldehyde concentration and were consistent with bimolecular kinetics (i.e., for propanal, *v*_on_ = *k*_on_[propanal]_tot_, where *k*_on_ is the apparent rate constant of adduct formation). The rates of hemithioacetal dissociation (*v*_off_) were independent of propanal concentration, consistent with a unimolecular step (i.e., *v*_off_ = *k*_off_, where *k*_off_ is the rate constant of adduct dissociation) (Fig. 1d and Supplementary Figs. 2-10). Both the association and dissociation reactions were pH-dependent and single-channel recordings were performed at pH 6.8. At this pH value, frequent events occurred at µM-mM analyte concentrations (e.g., ∼500 events within 10 min for 3 mM butanal), and the lifetimes of the hemithioacetal adducts were long to allow accurate aldehyde identification from I_res%_ values (e.g., ∼130 ms for butanal).

**Fig. 2:**
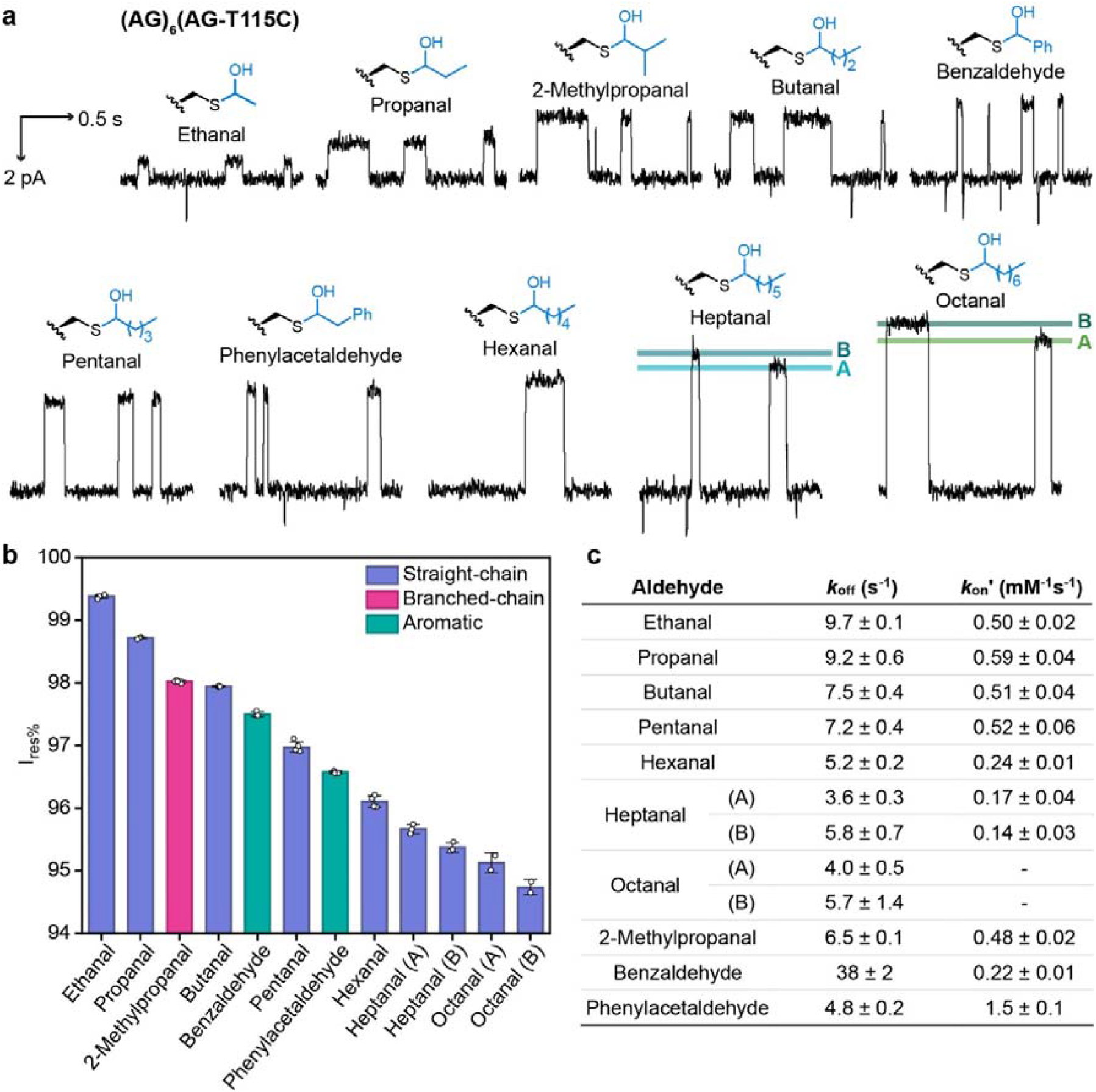
Thiol-aldehyde chemistry within the (AG)_6_(AG-T115C) nanopore. **a**, Aldehydes reacted with the cysteine at position 115 to generate hemithioacetal adducts reversibly. Diastereomeric adducts were resolved for heptanal and octanal. Diastereomer A is arbitrarily assigned with a higher I_res%_ than diastereomer B. Recording conditions: −50 mV (trans) with 2 M KCl, 200 mM PIPES and 20 μM EDTA at pH 6.8. Aldehyde concentrations (trans): 5.8 mM ethanal, 5.8 mM propanal, 6.1 mM 2-methylpropanal, 5.8 mM butanal, 5.7 mM benzaldehyde, 5.5 mM pentanal, 5.5 mM phenylacetaldehyde, 2.7 mM hexanal, 1.5 mM heptanal, 1.3 mM octanal. Signals were low-pass filtered at 10 kHz and sampled at 50[kHz. Traces were further filtered at 100 Hz for display. **b**, I_res%_ values for the hemithioacetal adducts. **c**, Rate constants for adduct formation corrected for hydration (*k*_on_□) and dissociation (*k*_off_). Errors are standard deviations across 3 different nanopores. Corrected rate constants for adduct formation were obtained from apparent rate constants (*k*_on_) by using the equation: *k*_on_□ = *k*_on_ × (1 + K_hyd_). Rates of adduct formation for octanal were not reported due to the poor solubility of octanal in recording solutions, precluding accurate determinations.

In total, 10 different aldehydes were characterized with the (AG)_6_(AG-T115C) nanopore (Fig. 2a and Supplementary Table 1), ranging from straight-chain to branched-chain to aromatic aldehydes. We demonstrated single CH_2_ resolution in distinguishing straight-chain aldehydes (Fig. 2b and Supplementary Table 1). From ethanal to octanal, each additional CH_2_ group reduced the I_res%_ value by ∼0.7 %. Interestingly, our approach was able to distinguish between diastereomeric hemithioacetal adducts with opposite chirality at the Ca position for both heptanal and octanal (ΔI_res%_ = 0.29 ± 0.03 % for heptanal (N = 3 pores, >100 events for each diastereomer); 0.39 ± 0.05 % for octanal (N = 2 pores, >40 events for each diastereomer)). We arbitrarily assigned adducts with larger I_res%_ as diastereomers A and those with smaller I_res%_ as diastereomers B (i.e., I_res%,A_ > I_res%,B_). For shorter straight-chain aldehydes, diastereomeric adducts were not clearly separated by (AG)_6_(AG-T115C) with ΔI_res%_ <0.2 %. Aromatic aldehydes produced higher I_res%_ values than saturated aliphatic aldehydes with similar masses, i.e., I_res%_ = 97.5 ± 0.1 % for benzaldehyde (N = 3 pores, >300 events) and 96.1 ± 0.1 % for hexanal (N = 3 pores, >400 events); I_res%_ = 96.6 ± 0.1 % for phenylacetaldehyde (N = 4 pores, >400 events) and 95.7 ± 0.1 % and 95.4 ± 0.1 % for heptanal (N = 3 pores, >100 events for each diastereomer). We speculate that the open-chain aldehydes extend away from the protein wall, creating larger steric blockades than the corresponding cyclic aromatic aldehydes. Branched-chain aldehydes and their straight-chain isomers produced similar I_res%_ values, i.e., I_res%_ = 98.1 ± 0.1 % for 2-methylpropanal (N = 5 pores, >500 events) and 97.9 ± 0.1 % for butanal (N = 4 pores, >400 events), with ΔI_res%_ ∼0.2 % for 2-methylpropanal and butanal simultaneously recorded with the same channel (N = 1 pore, >600 events).

### Simultaneous detection of aldehydes facilitated ratiometric profiling

To enable quantitative aldehyde detection, kinetic analysis of the formation and dissociation of hemithioacetal adducts was carried out. The solubility of each aldehyde in the recording solution was measured by NMR in the presence of internal standards (5 mM calcium formate and 5 mM maleic acid). Aldehydes formed hydrates to varying extents in aqueous solutions, and the hydrated forms were unreactive towards thiols and thus not detected by single-channel recording (Fig. 1b). We determined the hydration equilibrium constants (K_hyd_) by ^1^H NMR under electrical recording conditions (Supplementary Table 2 and Supplementary Fig. 11) and derived the corrected rate constants of adduct formation (*k*_on_□) by using the equations:

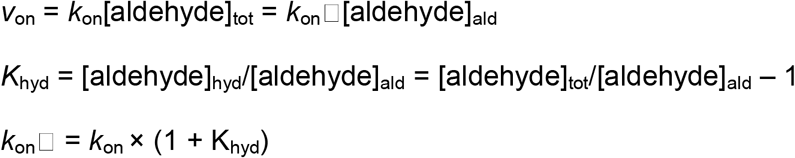

*v*_on_, rate of hemithioacetal formation; *k*_on_, apparent rate constant of hemithioacetal formation; *k*_on_□, corrected rate constant of hemithioacetal formation; *K*_hyd_, hydration equilibrium constant; [aldehyde]_tot_, total concentration of aldehyde; [aldehyde]_hyd_, concentration of aldehyde in the hydrate form; [aldehyde]_ald_, concentration of free aldehyde.

For the bimolecular formation of hemithioacetal, *k*_on_□ remained at ∼0.5-0.6 mM^-1^s^-1^ for ethanal, propanal, butanal, and pentanal but dropped to ∼0.3 mM^-1^s^-1^ for hexanal and heptanal. While the energy difference is small, these observations could reflect the steric hindrance for the thiolate to approach the carbonyl group along the Bürgi-Dunitz angle, caused by the conformationally labile alkyl chains.^24^ As rates of adduct dissociation were independent of aldehyde concentration, no correction for hydration was required. For the unimolecular dissociation of hemithioacetals formed with straight-chain aldehydes, *k*_off_ remained within an order of magnitude, gradually decreasing as the chain length increases from 9.7 s^-1^ for ethanal to ∼5 s^-1^ for hexanal, heptanal and octanal (Fig. 2c). This could be attributed to the positive inductive effect of the increasing alkyl chain length.

Simultaneous detection of multiple aldehydes with a single nanopore was demonstrated with a mixture of 7 straight-chain aldehydes (i.e., ethanal to octanal) (Fig. 3a). The consistent current signatures for individual aldehydes and the clear separation between them enabled automated assignment of events by using machine learning (Fig. 3b). Almost 1000 individual events per aldehyde were collected from separate traces of each analyte (Supplementary Fig. 12). A stratified random split was performed, leaving 30% of the events as the test set. Three features were extracted to characterize each event: I_res%_, event duration, and the root-mean-squared noise of the event (See Supplementary Section 5). The random forest model achieved the highest accuracy of 98% on both the training and test sets, which was calculated as the fraction of correctly classified events in the dataset (Fig. 3b and Supplementary 13). Further, we demonstrated ratiometric determination of aldehydes in mixtures with pentanal and butanal, for which the detection frequencies reflected their varied concentration ratios (Fig. 3c). For disease diagnosis and product quality control applications, ratiometric profiles are expected to be the most informative readout.

**Fig. 3:**
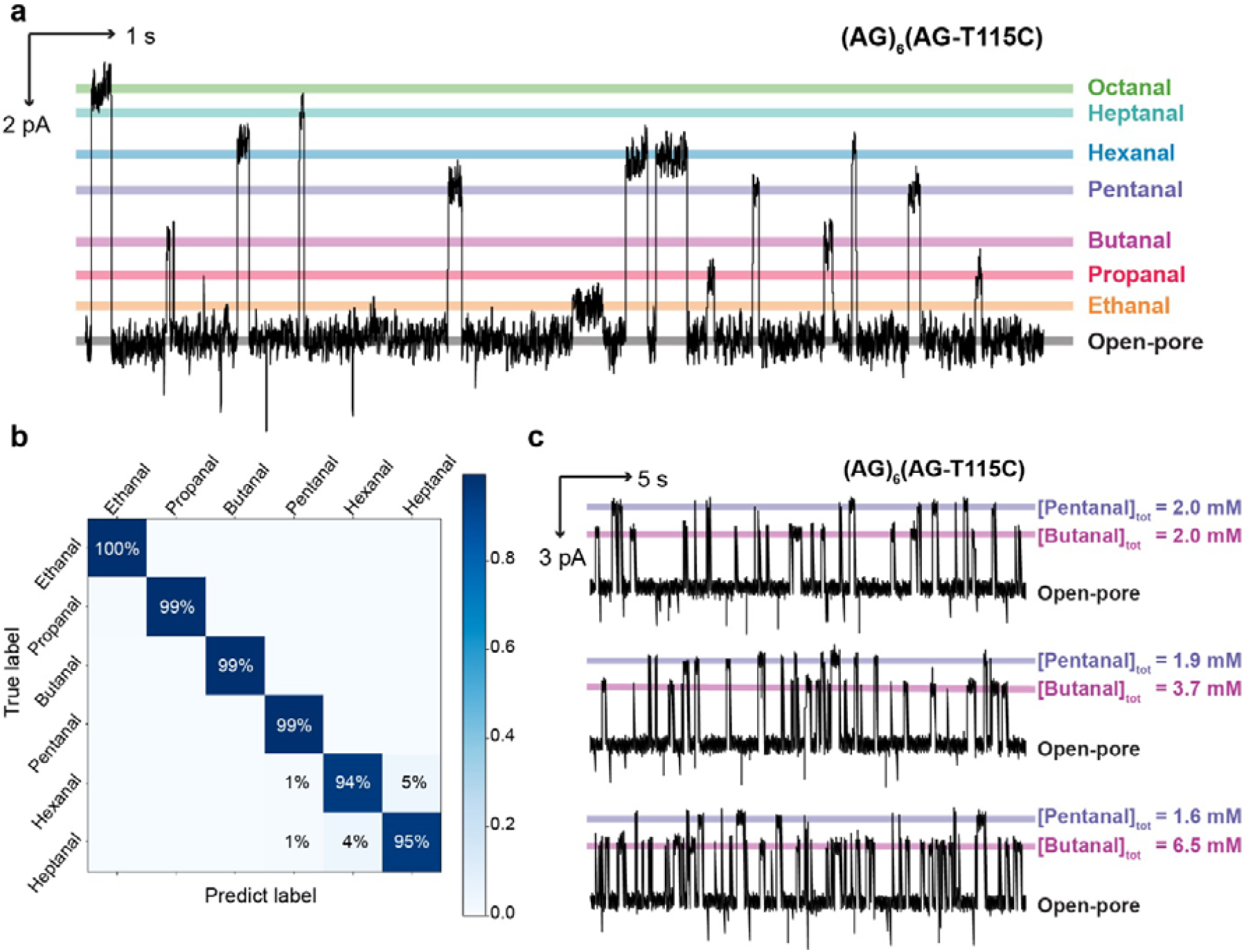
Single-molecule profiling of aldehyde mixtures. **a**, Simultaneous detection of 7 straight-chain aldehydes at the single-molecule level. Total aldehyde concentrations (trans): 0.7 mM ethanal, 0.7 mM propanal, 0.7 mM butanal, 0.5 mM pentanal, 0.6 mM hexanal, 0.5 mM heptanal, and 0.3 mM octanal. **b**, Performance of the Random Forest model on the test set is shown as a confusion matrix. The accuracy on the test set is 0.98, calculated as the ratio of correctly classified datapoints over all datapoints in the dataset. **c**, Single-channel recordings of pentanal and butanal at various ratios of total concentrations ([aldehyde]_tot_) as noted on the panel. Recording conditions: −50 mV (trans) with 2 M KCl, 200 mM PIPES and 20 μM EDTA at pH 6.8. Signals were low-pass filtered at 10 kHz and sampled at 50□kHz. Traces were further filtered at 100 Hz for display.

### Rational nanopore engineering for diastereomer and structural isomer resolution

Challenged by the similar current signatures observed with aldehyde structural isomers (e.g., butanal and 2-methylpropanal) with the (AG)_6_(AG-T115C) nanopore, rational nanopore engineering was undertaken. We hypothesised that current level resolution of diastereomeric hemithioacetal adducts would provide an additional layer of information to aid structural isomer resolution. To this end, we designed two additional nanopores in which the reactive cysteine residue was positioned within a narrower, and hence more sensitive, region of the nanopore β barrel (Fig. 4a). In the (MK)_6_(MK-T115C) nanopore (MK = WT-K8A-K131G-K147G), methionine residues were reintroduced at position 113 to reduce the internal diameter near the sensing site. In the (AG)_6_(AG-G137C) nanopore, the cysteine residue was moved to position 137, which was in close proximity to the narrowest region of the nanopore bearing an AG background (Supplementary Fig. 14). Three asparagine-to-alanine mutations around the sensing site were further introduced in to reduce sterics and promote diastereomeric interactions of hemithioacetal adducts with the local protein environment, leading to the (AG)_6_(AG-G137C-Ala3) nanopore (AG-G137C-Ala = AG-G137C-N139A-N121A-N123A) (Fig. 4a).

**Fig. 4:**
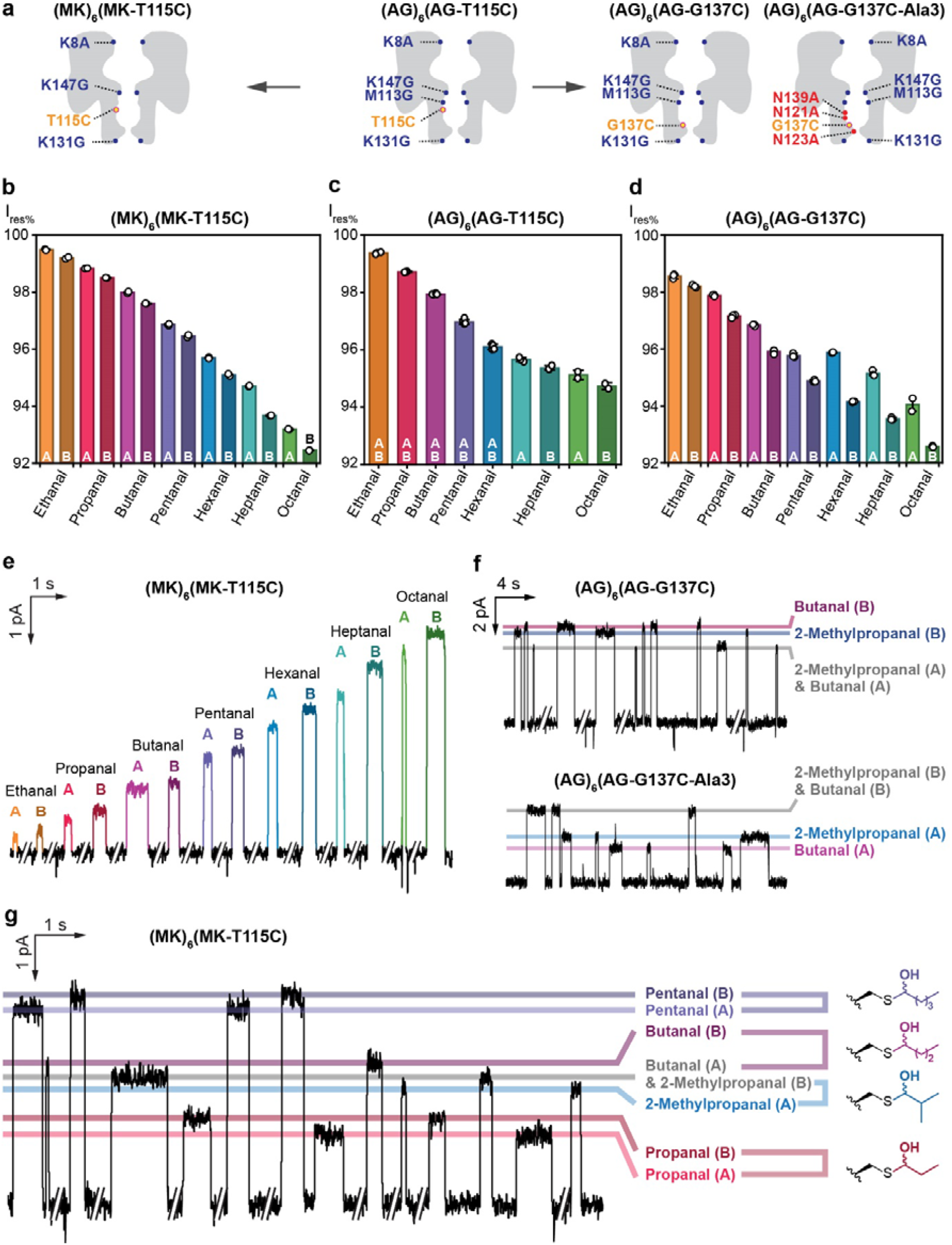
Diastereomer discrimination in engineered nanopores facilitated structural isomer resolution. **a**, Rational engineering of nanopores to enhance diastereomer discrimination through two parallel strategies: Left: narrowing the pore diameter around the sensing group (yellow); Right: moving the sensing group to the narrowest internal site. Mutations are highlighted relative to the WT background in all subunits (blue) or in the cysteine-bearing subunit (red). **b-c**, Within the (MK)_6_(MK-T115C) nanopore, diastereomeric adducts were resolved for all straight-chain aldehydes tested. Diastereomer A was arbitrarily assigned with a higher I_res%_ than diastereomer B. **d**, Within the (AG)_6_(AG-T115C) nanopore, diastereomeric adducts were resolved for only heptanal and octanal. **e**, Within the (AG)_6_(AG-G137C) nanopore, diastereomeric adducts were resolved for all straight-chain aldehydes tested. **f**, A pair of chain isomers, butanal and 2-methylpropanal, were distinguished by diastereomers B in the (AG)_6_(AG-G137C) nanopore or by diastereomers A in the (AG)_6_(AG-G137C-Ala3) nanopore. **g**, Propanal, butanal, 2-methylpropanal, and pentanal were detected within a single (MK)_6_(MK-T115C) pore. Butanal and 2-methylpropanal were distinguished by the diastereomer A of 2-methylpropanal and the diastereomer B of butanal. Recording conditions: −50 mV (trans) with 2 M KCl, 200 mM PIPES and 20 μM EDTA at pH 6.8. Signals were low-pass filtered at 10 kHz and sampled at 50□kHz. Traces were further filtered at 50 Hz for display.

The I_res%_ of 7 different straight-chain aldehydes (i.e., ethanal to octanal) were characterized with the (MK)_6_(MK-T115C) and (AG)_6_(AG-G137C) nanopores (Fig. 4b-c and Supplementary Tables 4-5). Improved diastereomer discrimination was observed in both nanopores: diastereomeric hemithioacetal adducts from ethanal to octanal were easily separable in both nanopores: 0.3 % < ΔI_res%_ < 1.0 %, in the (MK)_6_(MK-T115C) nanopore (Supplementary Table 4) and 0.4 % < ΔI_res%_ < 1.6 %, in the (AG)_6_(AG-G137C) nanopore (Supplementary Table 5). Good current level separation was achieved across all diastereomeric adducts in the (MK)_6_(MK-T115C) nanoreactor, whereas some overlap was seen in the (AG)_6_(AG-G137C) nanoreactor (e.g., I_res%_ = 95.8 ± 0.1 % for diastereomer A of pentanal and 95.9 ± 0.1 % for diastereomer A of hexanal).

Current level resolution of diastereomeric hemithioacetal adducts subsequently allowed for structural isomer resolution. In the (AG)_6_(AG-G137C) nanopore, butanal and 2-methylpropanal could be distinguished based on diastereomers B (i.e., ΔI_res%_ of 0.25 ± 0.02 % for diastereomers B of butanal and 2-methylpropanal, N = 3 pores, >35 events for each aldehyde) (Supplementary Fig. 15). In contrast, within the (AG)_6_(AG-G137C-Ala3) nanopore, diastereomer pairs were better resolved for both butanal or 2-methylpropanal; distinct separation between diastereomers A enabled clear resolution of these chain isomers (i.e., ΔI_res%_ of 0.42 ± 0.03 % for diastereomers A of butanal and 2-methylpropanal, N = 4 pores, >60 events for each aldehyde). In the (MK)_6_(MK-T115C) nanopore, butanal and 2-methylpropanal could be distinguished using diastereomer A of 2-methylpropanal and diastereomer B for butanal (i.e., ΔI_res%_ of 0.43 ± 0.03 % for diastereomer B of butanal and diastereomer A of 2-methylpropanal, N = 2 pores, >50 events for each aldehyde). As a proof of concept, simultaneous detection of propanal, butanal, 2-methylpropanal and pentanal was demonstrated in the (MK)_6_(MK-T115C) nanopore (Fig. 4d). As the current level blockades for pentanal and propanal do not overlap with those of butanal and 2-methylpropanal, all four aldehydes could be differentiated from I_res%_ alone.

### Enzyme-assisted differential sensing of alcohols and aldehydes

Nanopore covalent sensing is inherently selective for a targeted class of analytes. For example, in a mixture of alcohols (1-pentanol, 1-hexanol and 1-heptanol) and aldehydes (propanal and butanal), only the aldehydes were picked up by the (AG)_6_(AG-T115C) nanopore sensor (Fig. 5a). After treatment with an engineered alcohol oxidase^25^, additional aldehyde species—pentanal, hexanal, and heptanal— were identified (Fig. 5b), indirectly revealing the presence of alcohols in the original sample. This simple functional group conversion step, coupled to nanopore sensing, therefore allowed for differential single-molecule detection of alcohols and aldehydes in a mixture—a powerful strategy worth further development.

**Fig. 5:**
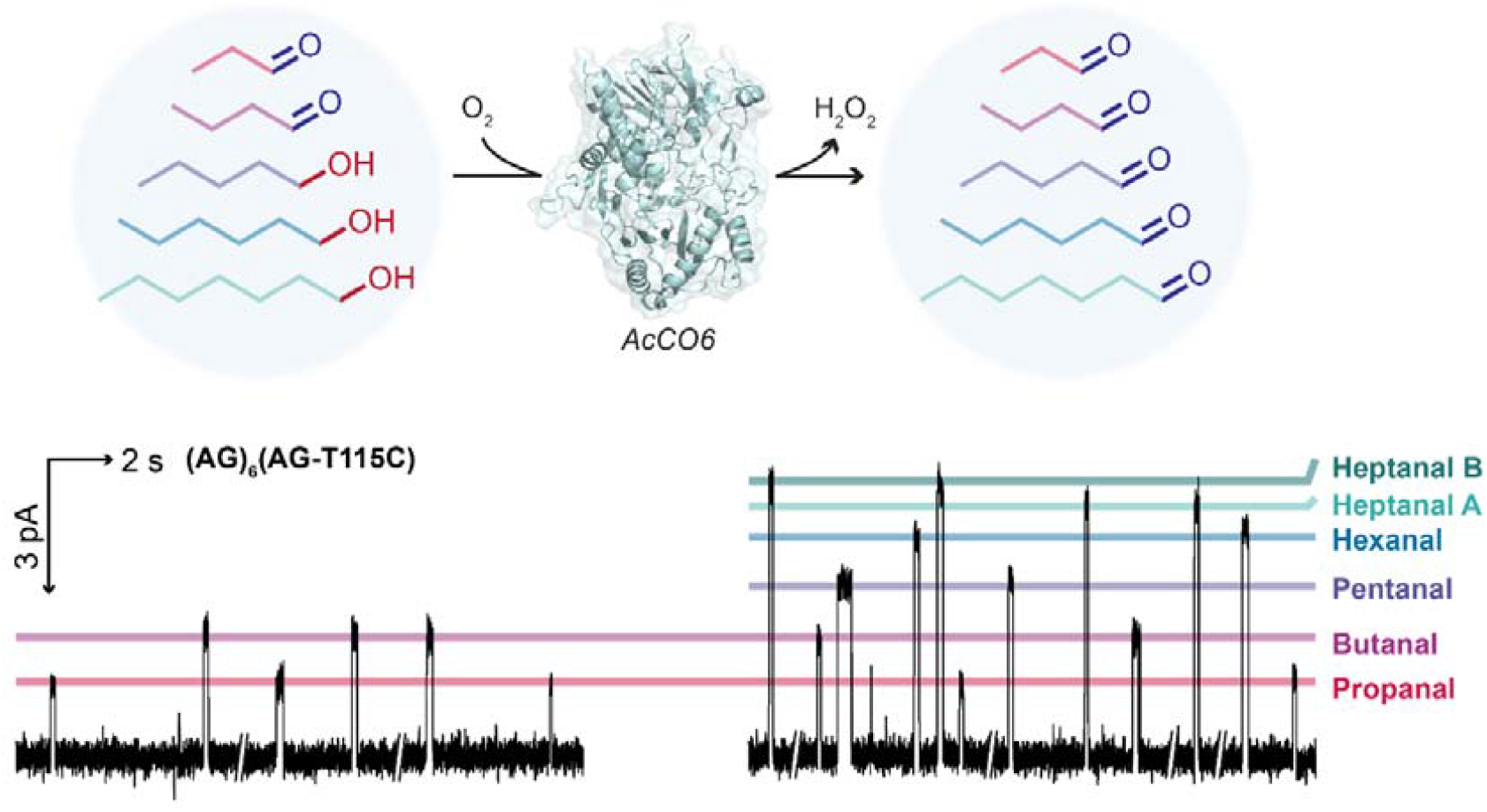
Enzyme-facilitated indirect alcohol detection. The (AG)_6_(AG-T115C) nanopore selectively detected aldehydes in an aldehyde-alcohol mixture (Left). After treatment with an engineered alcohol oxidase (AcCO6), additional aldehydes species—pentanal, hexanal, heptanal—were identified (Right), which indirectly confirmed the presence of the corresponding alcohols in the original sample. Recording conditions: −50 mV (trans) with 2 M KCl, 200 mM PIPES and 20 μM EDTA at pH 6.8. Signals were low-pass filtered at 10 kHz and sampled at 50□kHz. Traces were further filtered at 100 Hz for display.

## Conclusions

The present work exploited thiol-aldehyde chemistry, a previously underexplored dynamic covalent chemistry, within protein nanopores to target the sensing of aldehydes, a key class of chemicals ubiquitous in daily life. We achieved single-molecule identification of 10 straight-chain, branched-chain, and aromatic aldehydes, which represent potential biomarkers for lung cancer,^5^ Crohn’s disease,^6^ SARS-CoV-2,^8^ as well as atmospheric pollutants,^2^ and beverage impurities.^3^ Our approach resolved straight-chain aldehydes that differed by single CH_2_ groups, members of hemithioacetal diastereomeric pairs, and structural isomers—challenges often faced by conventional small-molecule detection methods. In particular, we systematically engineered the protein environment surrounding the covalent sensing site, achieving improved separation for diastereomeric adducts by promoting interactions with the pore interior. Simple engineering of the sensing region thus holds promise for improving small-molecule covalent detection with a given nanopore scaffold.

As a proof of concept, we established ratiometric profiles of mixed aldehydes at mM concentrations with >500 events collected in 10 min using a single nanopore. Given that common biological samples contain aldehydes at µM concentrations^26,27^, routine methods can be applied to capture and concentrate analytes.^28^ To analyze biological samples directly, the detection frequency could be further increased by >500,000-fold using nanopore mutants engineered with more than one covalent sensing sites (e.g., (AG-T115C)_7_), higher recoding pH values, elevated temperatures, and nanopore sensing devices containing arrays of pores (e.g. the PromethION device contains up to 2675 nanopores per flow cell, and 48 flow cells per machine). Based on our findings, we envision an accessible nanopore platform for rapid aldehyde detection, paving the way for applications in disease diagnosis, environmental monitoring, and the quality control of food and beverages.

Moving forward, the detection of other biologically relevant chemical classes can be explored by capitalizing the aldehyde-sensing system here. As central metabolites, aldehydes are generated by a wide range of enzymes from various functional groups, including carboxylic acids, primary alcohols, and primary amines^29^. Many of these convertible chemical classes are also potential disease biomarkers. For example, 1-pentanol, detected as 1-pentanal in this work, has been found in the exhaled breath by lung cancer patients but not healthy individuals^5^. While we have demonstrated alcohol sensing after enzymatic conversion to aldehydes, detecting other chemical classes requires careful identification of suitable enzymes, particularly with respect to substrate scope and catalytic efficiency. In the long term, by leveraging thiol-aldehyde sensing chemistry, we envision a versatile sensing workflow that employs a suite of aldehyde-converting reagents, which will produce informative single-molecule profiles of various chemical classes for nanopore diagnostics.

## Methods

### Preparation of nanopore sensors

Nanopore heteroheptamers were prepared according to the procedure below, which is a modification of a method previously reported.^30^ αHL monomers were prepared with an *E. coli in vitro* transcription and translation (IVTT) system (*E. coli* T7 S30 Extract System for Circular DNA, Cat #L1130, Promega). A standard reaction comprised: DNA plasmid mixture (<4 µg, plasmids encoding cysteine-free and cysteine-containing subunits were in an 8:1 ratio), amino acid mixture without methionine (5 µL, as supplied in the kit), S30 premix without amino acids (20 µL, as supplied in the kit), [^35^S]methionine (2 µL, 1200 Ci/mmol, 15 mCi/mL, MP Biomedicals), T7 S30 extract for circular DNA (15 µL, as supplied in the kit), and rabbit red blood cell membranes (2 µL, ∼ 1 mg protein/mL). The reaction mixture was incubated at 37°C for 2 h.

αHL heptamers containing different numbers of mutant subunits were separated in the gel based on their electrophoretic mobilities which were determined by the number of octa-aspartate (D8) tails present (i.e., each cysteine-bearing mutant subunit contained a D8 tail). Hence, the top band corresponded to homoheptamers bearing no octa-aspartate tail (i.e., AG-αHL)_7_, the second band corresponded to heteroheptamers bearing a single octa-aspartate tail (i.e., (mutant-D8)_1_(AG-αHL)_6_) and so on, with consecutive bands having an aspartate-tail-free subunit replaced with a mutant subunit bearing an octa-aspartate tail. In this work, the desired protein pore containing a single cysteine residue was extracted from the second band from the top.

### Single-channel electrical recordings

Single-channel recordings were carried out in a planar bilayer apparatus as previously described.^31^ A single αHL pore was allowed to insert into the bilayer. Aldehyde substrates were introduced from the trans compartment. Experiments were conducted using recording buffer containing 2 M KCl, 200 mM PIPES and 20 μM EDTA titrated to pH 6.8. Aldehyde solutions were prepared with recording buffer and titrated to pH 6.8. Single-channel recordings were conducted with a coverslip placed atop the recording chamber.

Ionic currents were recorded by using a patch clamp amplifier (Axopatch 200B, Axon Instruments), and filtered with a low-pass Bessel filter (80 dB/decade) with a corner frequency of 10 kHz. Signals were digitized with a Digidata 1320A digitizer (Molecular Devices) at an acquisition frequency of 50 kHz. The current traces were processed with Clampfit 10.7 (Molecular Devices). Current traces were idealized by using Clampfit 10.7 (Molecular Devices). The idealized data were analyzed with QuB 2.0 software (www.qub.buffalo.edu).^32^ Dwell time analysis and rate constant determinations were performed by using the maximum interval likelihood (MIL) algorithm of QuB.^33^

## Acknowledgements

This research was supported by the Bill & Melinda Gates Foundation, a European Research Council Advanced Grant (SYNTISU), a European Research Council Starting Grant (NANOPRO). We thank Steven Barry at the Physical and Theoretical Chemistry Laboratory, University of Oxford for fabricating the recording chamber.

## Author information

### Contributions

H.B. and Y.Q. conceived and supervised the project. L.E.M. and Z.H.L. prepared the proteins and conducted the single-channel recording experiments. Z.B. prepared the script for machine learning and performed all molecular dynamics simulations. Y.Y. prepared the AcCO6 proteins. L.E.M., Z.H.L., Z.B. and Y.Q. performed the data analysis. L.E.M., Z.H.L., Y.Y., Z.B., H.B. and Y.Q. wrote the manuscript.

## Ethics declarations

### Competing interests

H.B. is the founder of, a consultant for and a shareholder of Oxford Nanopore Technologies, a company engaged in the development of nanopore sensing and sequencing technologies. L.E.M., Z.H.L., H.B. and Y.Q. have filed patents describing the engineered nanopores and their applications in small-molecule covalent sensing.

Z.B. and Y.Y. are listed as contributors to the IP.

