## Supplementary Information for "Targeted, high-resolution sensing of volatile organic compounds by covalent nanopore detection"

### **1 Rationale for using α-hemolysin nanopores with AG background**

Background reactions were observed between benzaldehyde (20 mM, trans) and the wild-type α-hemolysin (αHL) pore ((WT-αHL)_7_), resulting in current levels A, B, C, and D in addition to the open pore level (level P) (Supplementary Fig. 1). These events were eliminated after replacing lysine residues at the cis entrance (Lys-8) and inward-facing lysine residues within the β barrel (Lys-131 and Lys-147). Additionally, the inward-facing methionine residues within the β barrel (Met-113) were replaced to eliminate any potential metal-binding sites within the nanopore.^1^ These mutations (i.e., K8A, M113G, K131G and K147G) were collectively called AG mutations. Further studies to identify the adducts producing the undesirable current levels were not conducted.

No detectable reactivity was observed with the (AG)_7_ homoheptamer (i.e., AG = WT-K8A-M113G-K131G-K147G) when up to 30 mM of benzaldehyde was introduced into the trans compartment. Hence, events observed when the (AG)_6_(AG-T115C) nanopore was used were attributed to the presence of the Cys-115 residue, specifically reversible hemithioacetal formation.

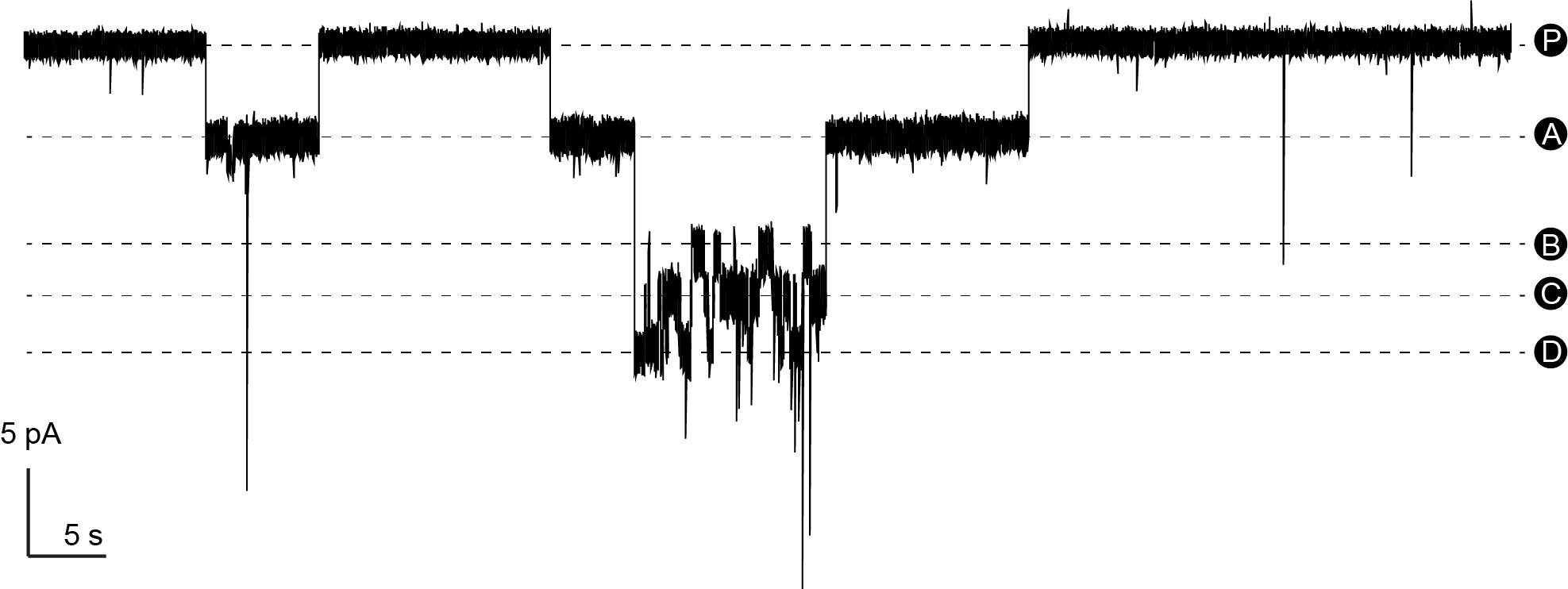
**Supplementary Fig. 1: Background reactions between benzaldehyde and the WT αHL pore ((WT-αHL)_7_).** Levels A, B, C, and D, which occurred in addition to the open pore current level (level P), were attributed to benzaldehyde reacting with lysine residues lining the β-barrel and the *cis* entrance. Conditions: 20 mM benzaldehyde (trans), (WT-αHL)_7_ nanopore, 2 M KCl, 200 mM HEPBS (pH 8.5), 20 μM EDTA, recorded at +50 mV (trans) and 21 ± 1 °C. Signals were low-pass filtered at 10 kHz and sampled at 50 kHz.

### **2** **Concentration-dependence of hemithioacetal formation: Kinetic studies**

Single-channel recordings were carried out at -50 mV (trans) with 2 M KCl, 200 mM PIPES and 20 μM EDTA at pH 6.8. Signals were low-pass filtered at 10 kHz and sampled at 50 kHz. Traces were further filtered at 100 Hz for display. Errors are standard deviations of rates at each concentration.

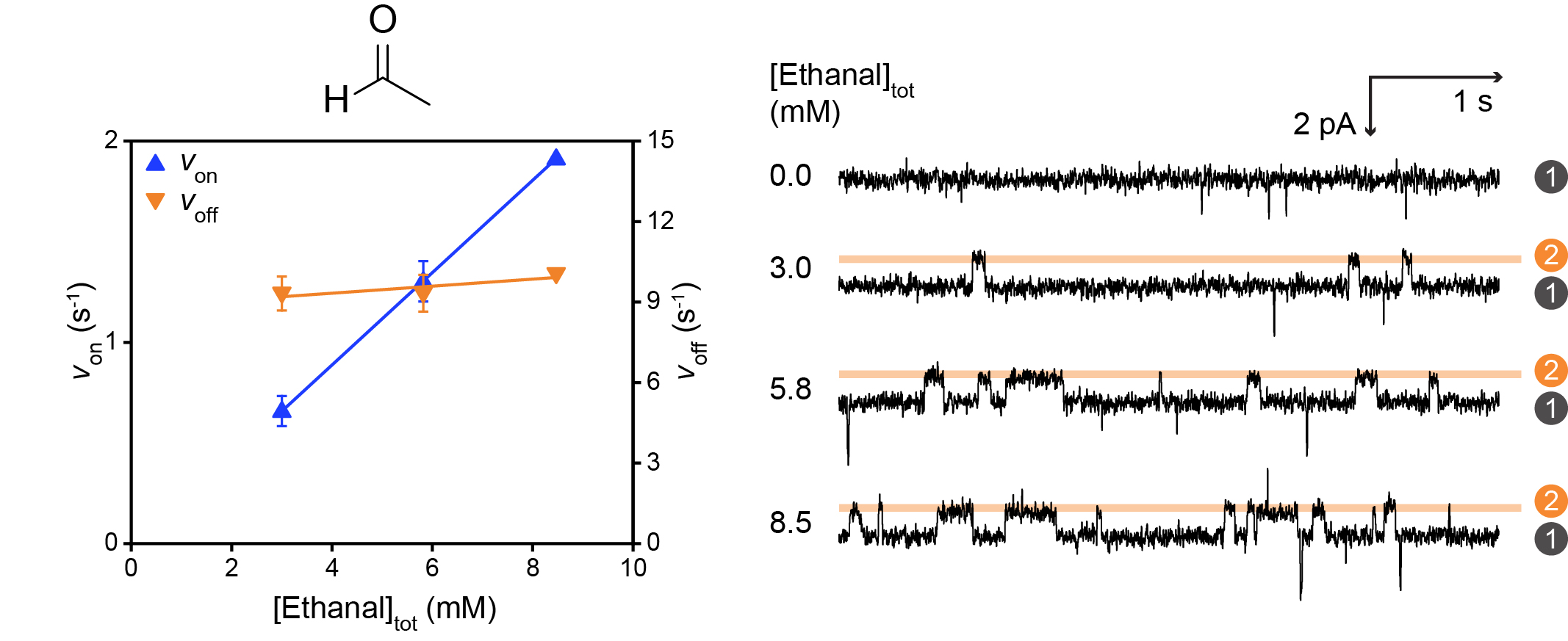

**Supplementary Fig. 2: Kinetic analysis of ethanal.** Left: Rates of adduct formation (*v*_on_) and dissociation (*v*_off_) plotted against total ethanal concentration. Errors are standard deviations of rates at each concentration. Right: Single-channel recordings with 0.0, 3.0, 5.8 and 8.5 mM total ethanal (trans). Current levels correspond to the hemithioacetal adducts (2) and the unoccupied nanopore (1).

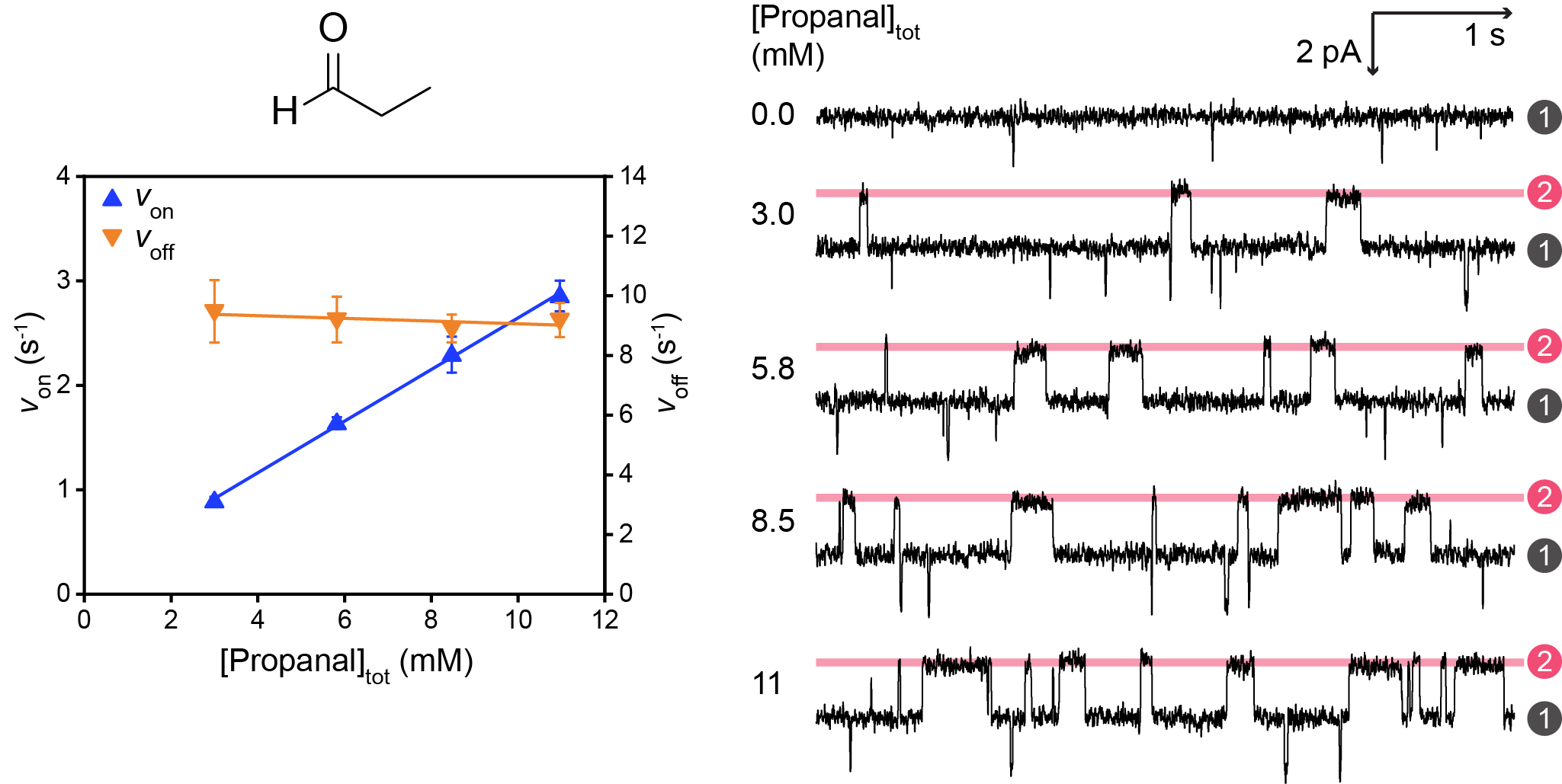

**Supplementary Fig. 3: Kinetic analysis of propanal.** Left: Rates of adduct formation (*v*_on_) and dissociation (*v*_off_) plotted against total propanal concentration. Errors are standard deviations of rates at each concentration. Right: Single-channel recordings with 0.0, 3.0, 5.8, 8.5 and 11 mM total propanal (trans). Current levels correspond to the hemithioacetal adducts (2) and the unoccupied nanopore (1).

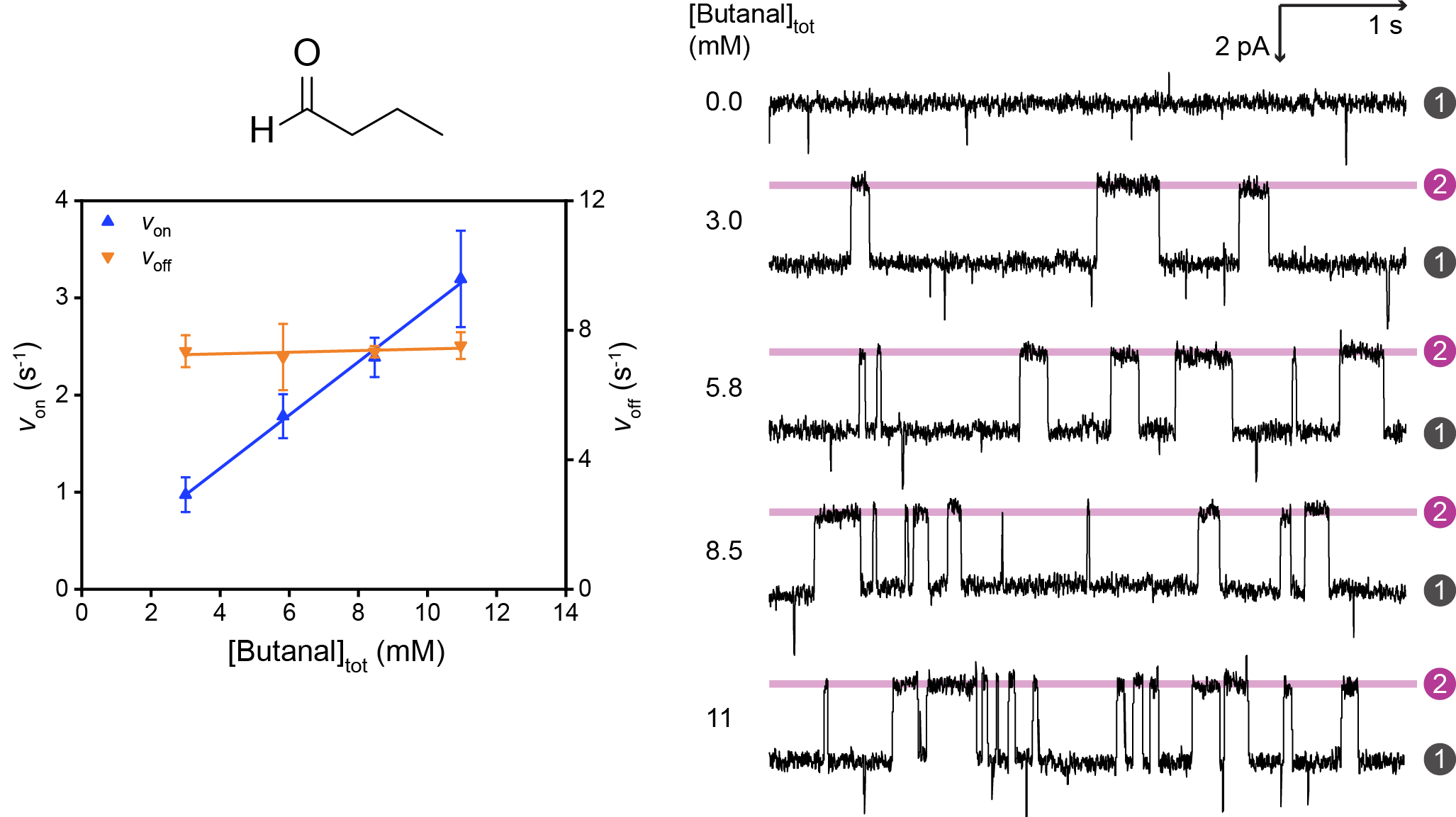

**Supplementary Fig. 4: Kinetic analysis of butanal.** Left: Rates of adduct formation (*v*_on_) and dissociation (*v*_off_) plotted against total butanal concentration. Errors are standard deviations of rates at each concentration. Right: Single-channel recordings with 0.0, 3.0, 5.8, 8.5 and 11 mM total butanal (trans). Current levels correspond to the hemithioacetal adducts (2) and the unoccupied nanopore (1).

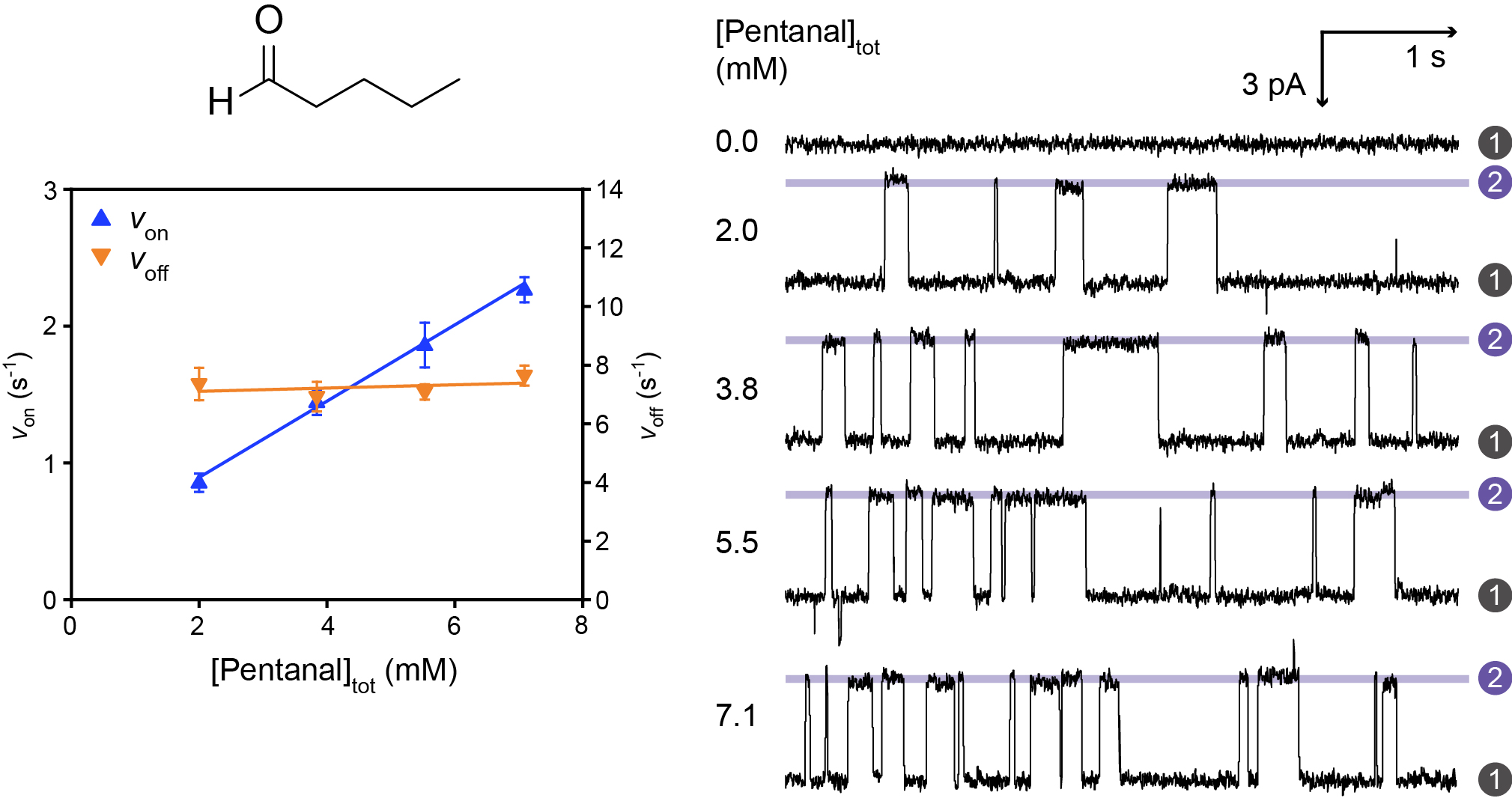

**Supplementary Fig. 5: Kinetic analysis of pentanal.** Left: Rates of adduct formation (*v*_on_) and dissociation (*v*_off_) plotted against total pentanal concentration. Errors are standard deviations of rates at each concentration. Right: Single-channel recordings with 0.0, 2.0, 3.8 and 5.5 mM total pentanal (trans). Current levels correspond to the hemithioacetal adducts (2) and the unoccupied nanopore (1).

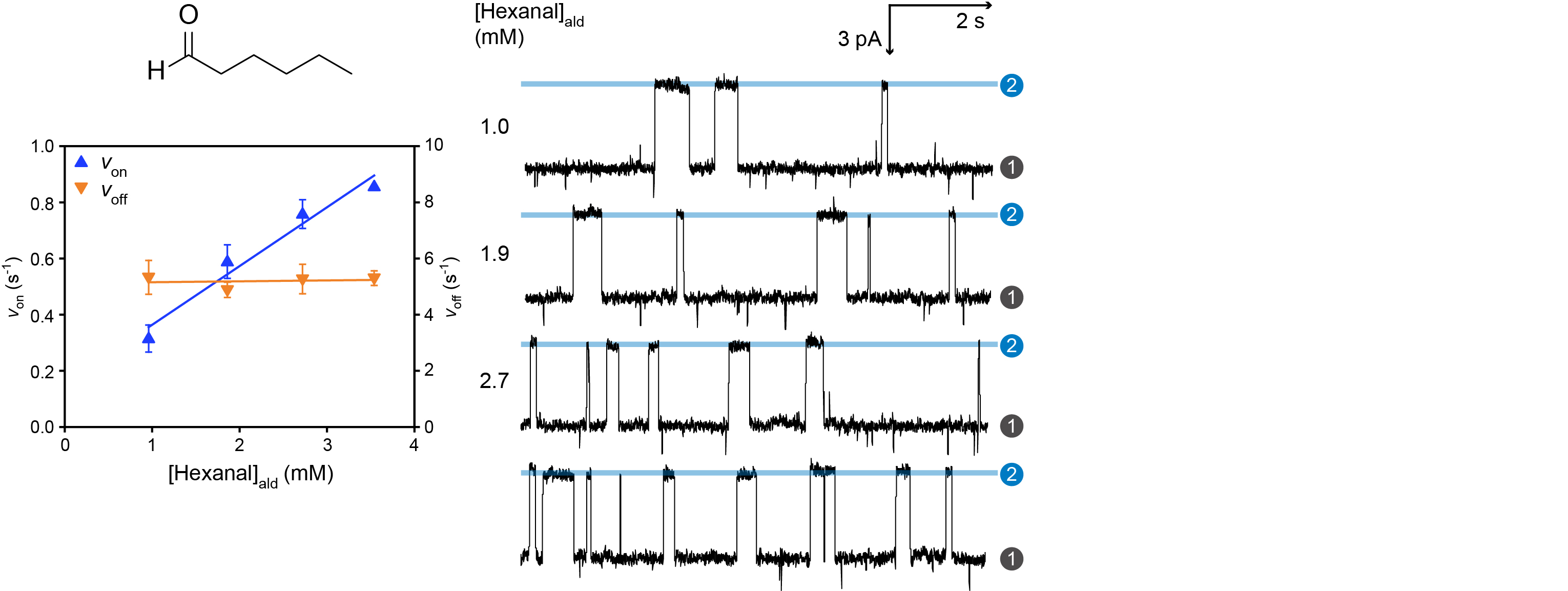

**Supplementary Fig. 6: Kinetic analysis of hexanal.** Left: Rates of adduct formation (*v*_on_) and dissociation (*v*_off_) plotted against the concentration of unhydrated hexanal (i.e., concentrations determined from ^1^H NMR, Supplementary Section 4). Errors are standard deviations of rates at each concentration. Right: Single-channel recordings with 1.0, 1.9 and 2.7 mM unhydrated hexanal (trans). Current levels correspond to the hemithioacetal adducts (2) and the unoccupied nanopore (1).

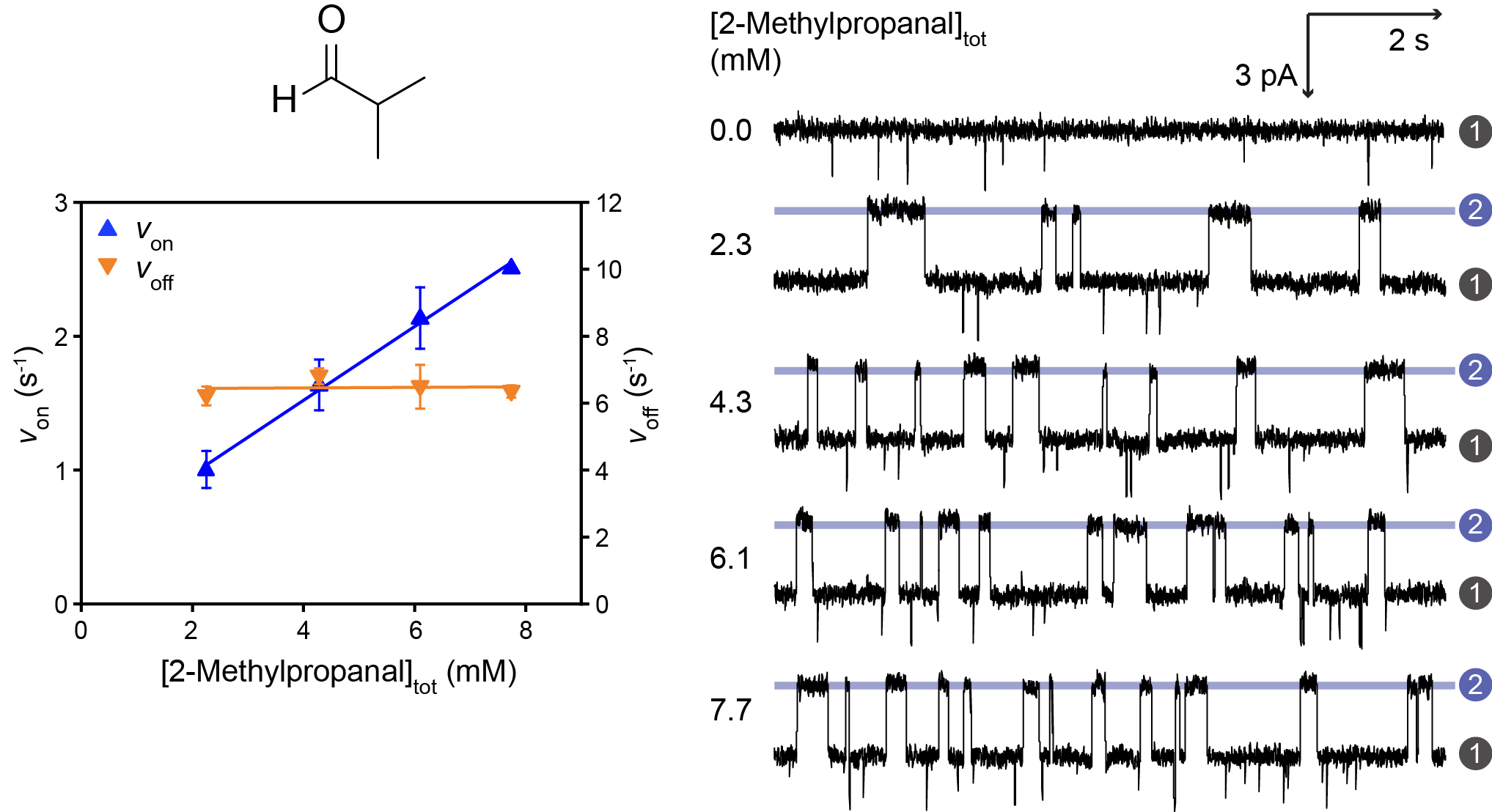

**Supplementary Fig. 7: Kinetic analysis of 2-methylpropanal.** Left: Rates of adduct formation (*v*_on_) and dissociation (*v*_off_) plotted against total 2-methylpropanal concentration. Errors are standard deviations of rates at each concentration. Right: Single-channel recordings with 0.0, 2.3, 4.3, 6.1 and 7.7 mM total 2-methylpropanal (trans). Current levels correspond to the hemithioacetal adducts (2) and the unoccupied nanopore (1).

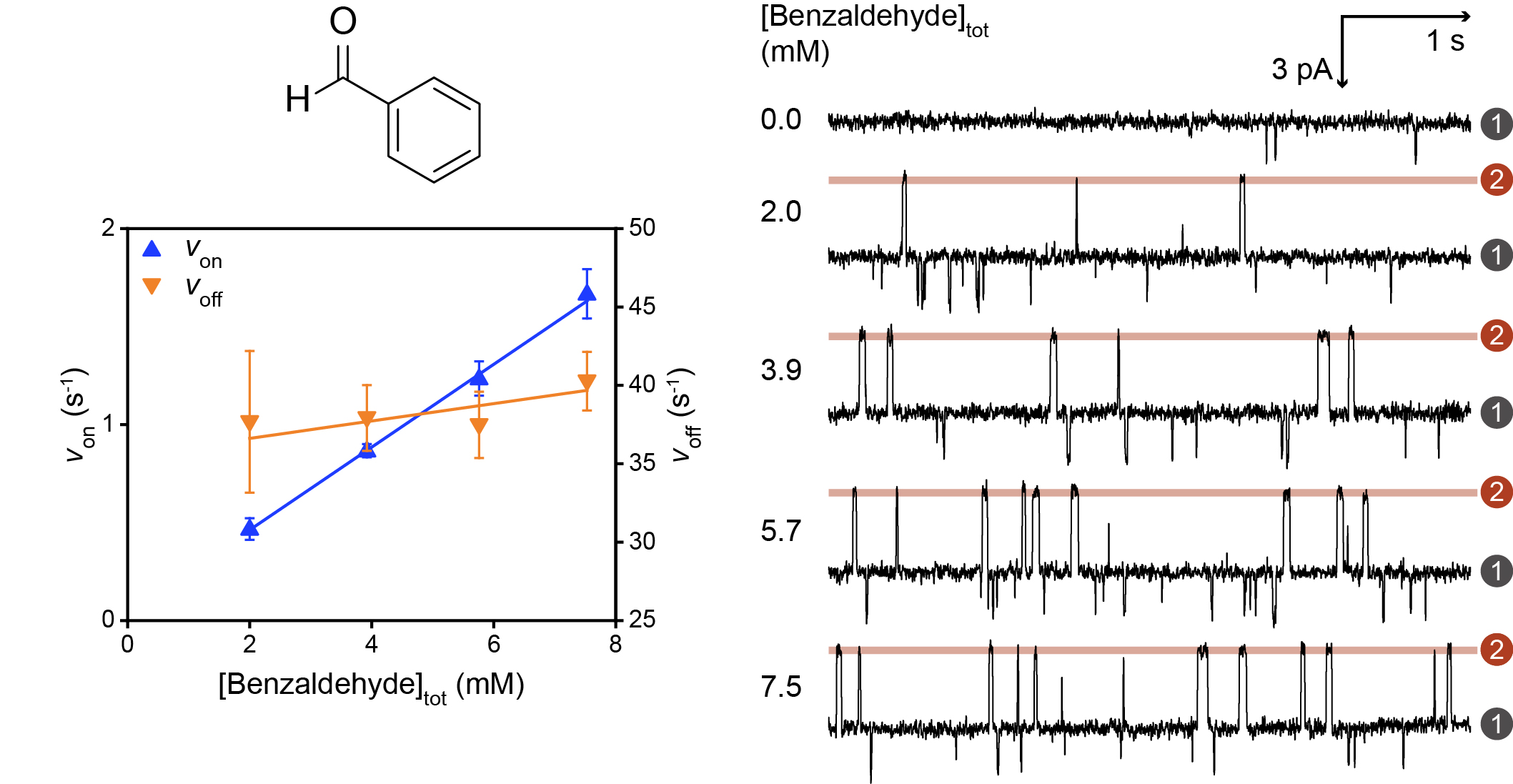

**Supplementary Fig. 8: Kinetic analysis of benzaldehyde.** Left: Rates of adduct formation (*v*_on_) and dissociation (*v*_off_) plotted against total benzaldehyde concentration. Errors are standard deviations of rates at each concentration. Right: Single-channel recordings with 0.0, 2.0, 3.9, 5.7 and 7.5 mM total benzaldehyde (trans). Current levels correspond to the hemithioacetal adducts (2) and the unoccupied nanopore (1).

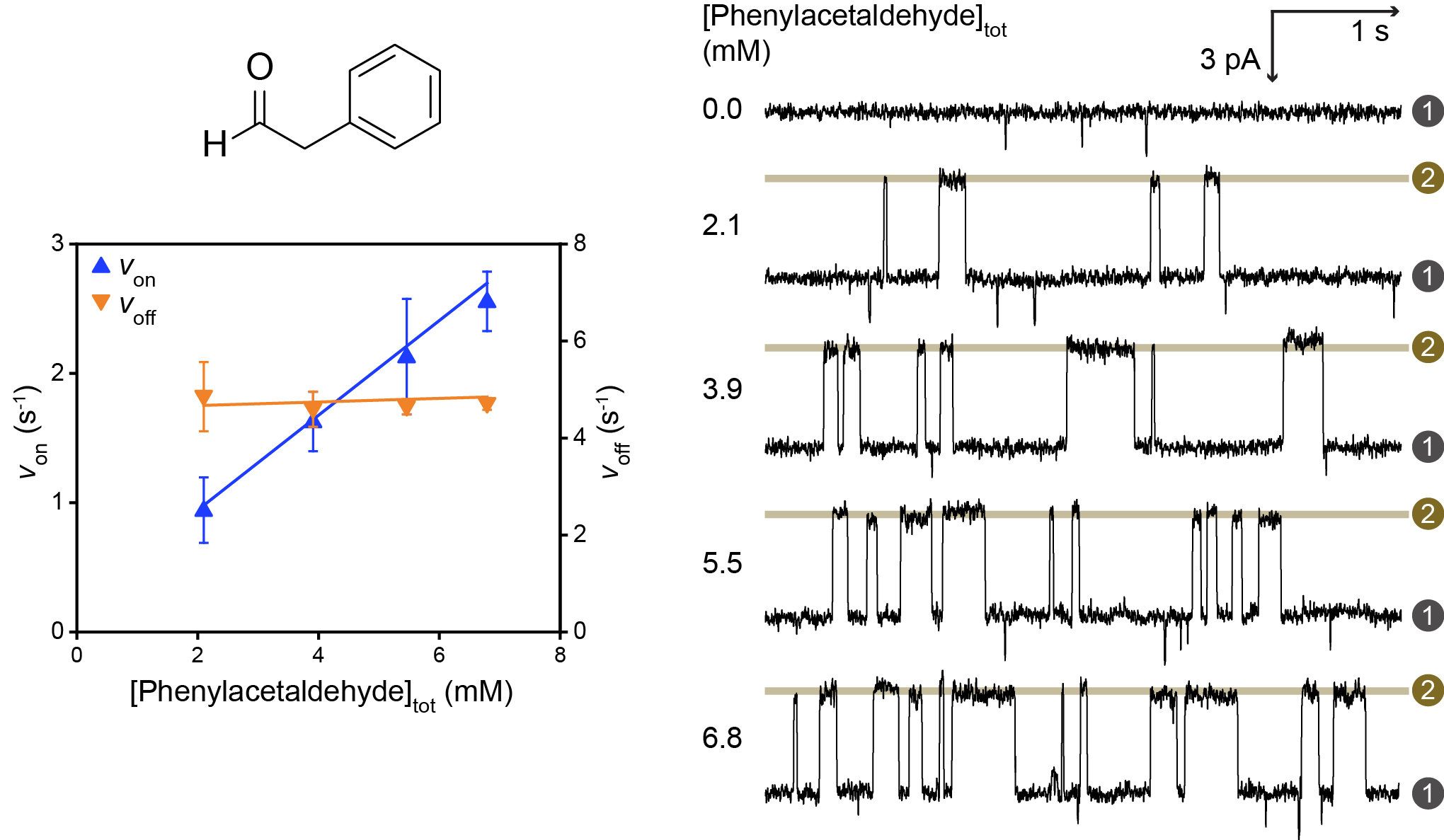

**Supplementary Fig. 9: Kinetic analysis of phenylacetaldehyde.** Left: Rates of adduct formation (*v*_on_) and dissociation (*v*_off_) plotted against total phenylacetaldehyde concentration. Errors are standard deviations of rates at each concentration. Right: Single-channel recordings with 0.0, 2.1, 3.9, 5.5 and 6.8 mM total phenylacetaldehyde (trans). Current levels correspond to the hemithioacetal adducts (2) and the unoccupied nanopore (1).

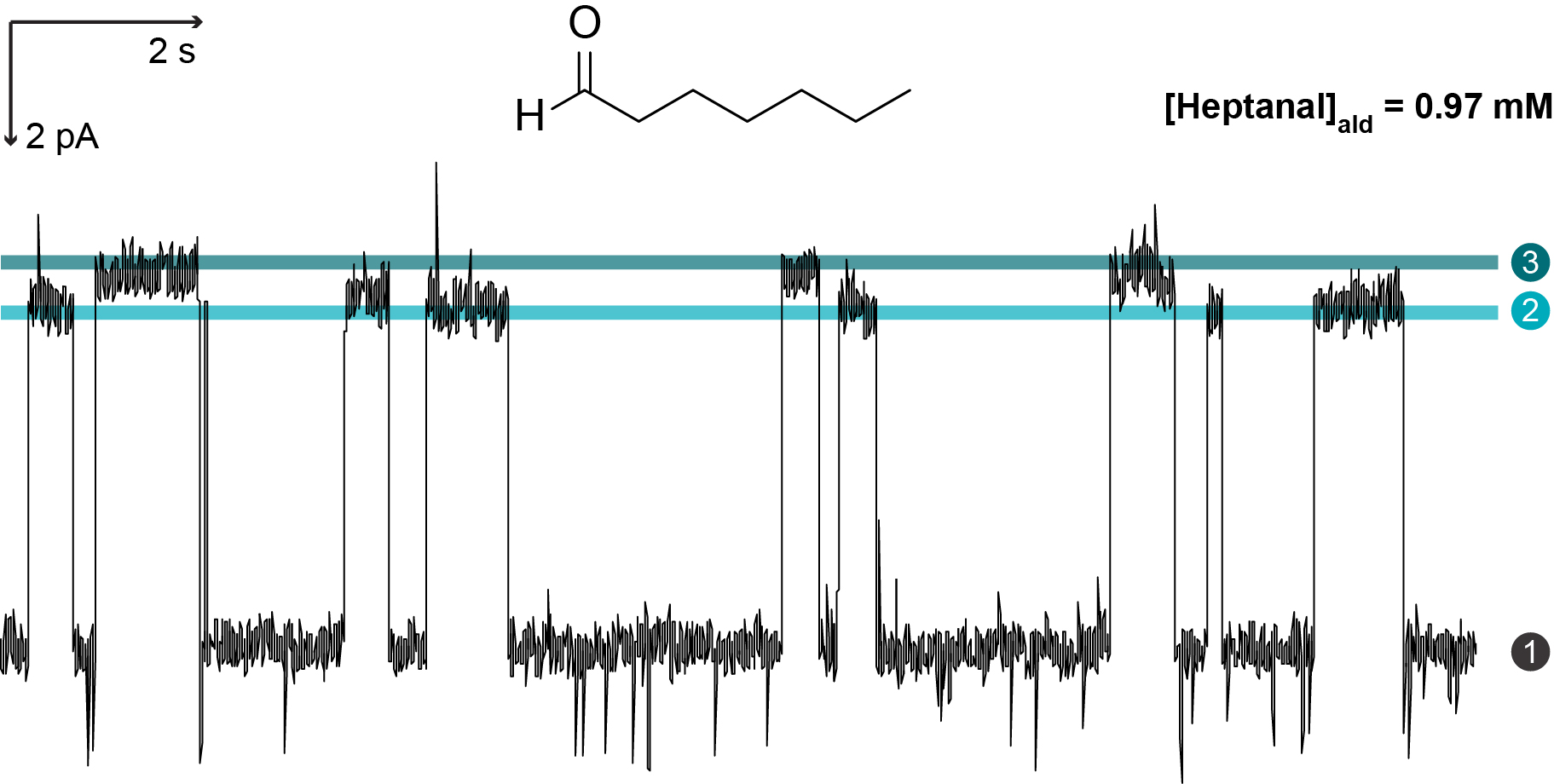

**Supplementary Fig. 10: Diastereomeric hemithioacetal adducts from heptanal have different conductance levels.** Single-channel recordings with 0.97 mM unhydrated heptanal (i.e., concentrations determined from ^1^H NMR, Supplementary Section 4). Current levels correspond to the two possible diastereomeric adducts, diastereomer A (2) and diastereomer B (3) and the unoccupied nanopore (1).

### **3 Kinetic rate constants for the reversible aldehyde-thiol chemistry and the percentage residual current levels for the hemithioacetal adducts**

| **Supplementary Table 1:** Apparent rate constants of formation (*k*_on_) and dissociation (*k*_off_), and percentage residual current levels (I_res%_) of hemithioacetal adducts.^[a,b,c]^ | | | | |
| --- | --- | --- | --- | --- |
| **Aldehyde** | | ***k*_on_ (mM^-1^s^-1^)** | ***k*_off_ (s^-1^)** | **I_res%_** |
| Ethanal | | 0.22 ± 0.01 | 9.7 ± 0.1 | 99.4 ± 0.1 |
| Propanal | | 0.25 ± 0.02 | 9.2 ± 0.6 | 98.7 ± 0.1 |
| Butanal | | 0.27 ± 0.02 | 7.5 ± 0.4 | 97.9 ± 0.1 |
| Pentanal | | 0.30 ± 0.03 | 7.2 ± 0.4 | 97.0 ± 0.1 |
| Hexanal | | 0.24 ± 0.01 | 5.2 ± 0.2 | 96.1 ± 0.1 |
| Heptanal | (A) | 0.17 ± 0.04 | 3.6 ± 0.3 | 95.7 ± 0.1 |
|  | (B) | 0.14 ± 0.03 | 5.8 ± 0.7 | 95.4 ± 0.1 |
| Octanal ^[d]^ | (A) | - | 4.0 ± 0.5 | 95.1 ± 0.1 |
|  | (B) | - | 5.7 ± 1.4 | 94.7 ± 0.1 |
| 2-Methylpropanal | | 0.27 ± 0.01 | 6.5 ± 0.1 | 98.1 ± 0.1 |
| Benzaldehyde | | 0.22 ± 0.01 | 38 ± 2 | 97.5 ± 0.1 |
| Phenylacetaldehyde | | 0.34 ± 0.01 | 4.8 ± 0.2 | 96.6 ± 0.1 |
| ^[a]^ Apparent rate constants have not been corrected to account for aldehyde hydration in solution.  ^[b]^ Conditions: (AG)_6_(AG-T115C) with 2 M KCl, 20 µM EDTA, 200 mM PIPES (pH 6.8), recorded at -50 mV (trans) and 22 ± 1 °C.  ^[c]^ Errors are standard deviations of the mean values from N > 3 nanopores, excluding octanal (N = 2 pores).  ^[d]^ The low solubility of octanal precluded the accurate determination of rate constants. | | | | |

### **4 Hydration constants and solubilities of aldehyde substrates**

Hydration constants of aldehyde substrates were determined by using ^1^H NMR. Solutions of ethanal, propanal, butanal, pentanal, 2-methylpropanal, benzaldehyde, and phenylacetaldehyde were prepared as follows. The aldehyde (50 mM for ethanal, propanal and butanal; 25 mM for pentanal and benzaldehyde; 22.5 mM for 2-methylpropanal; 15 mM for phenylacetaldehyde) was dissolved in a D_2_O solution of 200 mM PIPES and 2 M KCl at pH 6.4 (corresponding to a pD of 6.8),^2^ and left for 10 minutes to equilibrate before ^1^H NMR measurement. Hydration constants were calculated from the areas under the characteristic peaks of the aldehydes (∫_aldehyde_, aldehydic proton colored blue in Supplementary Fig. 11) and those of the corresponding hydrates (∫_hydrate_, gem-diol proton colored green in Supplementary Fig. 11). The aldehyde protons were found between chemical shifts δ 9.5-10.5, while the hydrate gem-diol protons were found between chemical shifts δ 4.5-5.5. An example spectrum for pentanal is shown (Supplementary Fig. 11).

For hexanal, heptanal and octanal, solutions were prepared as follows. In a D_2_O solution containing 200 mM PIPES and 2 M KCl at pH 6.4 (corresponding to a pD of 6.8), 5 mM calcium formate and 5 mM maleic acid were added as internal standards. To 1 mL of the D_2_O solution, 200 μL of aldehyde was added. The undissolved aldehyde formed a layer above the denser D_2_O layer. The biphasic mixture was stirred vigorously overnight for the D_2_O solution to become saturated with aldehyde. The saturated D_2_O layer was then used for ^1^H NMR measurement. Hydration constants were calculated from the areas under the characteristic peaks of aldehydes and those of the corresponding hydrates (Supplementary Table 2). Hydration constants for octanal were not determined due to the small area of the hydrate peak precluding accurate calculations. Concentrations of the unhydrated aldehydes in the saturated D_2_O layers were determined by comparing the areas of aldehyde signals to the areas of the internal standards (Supplementary Table 3).

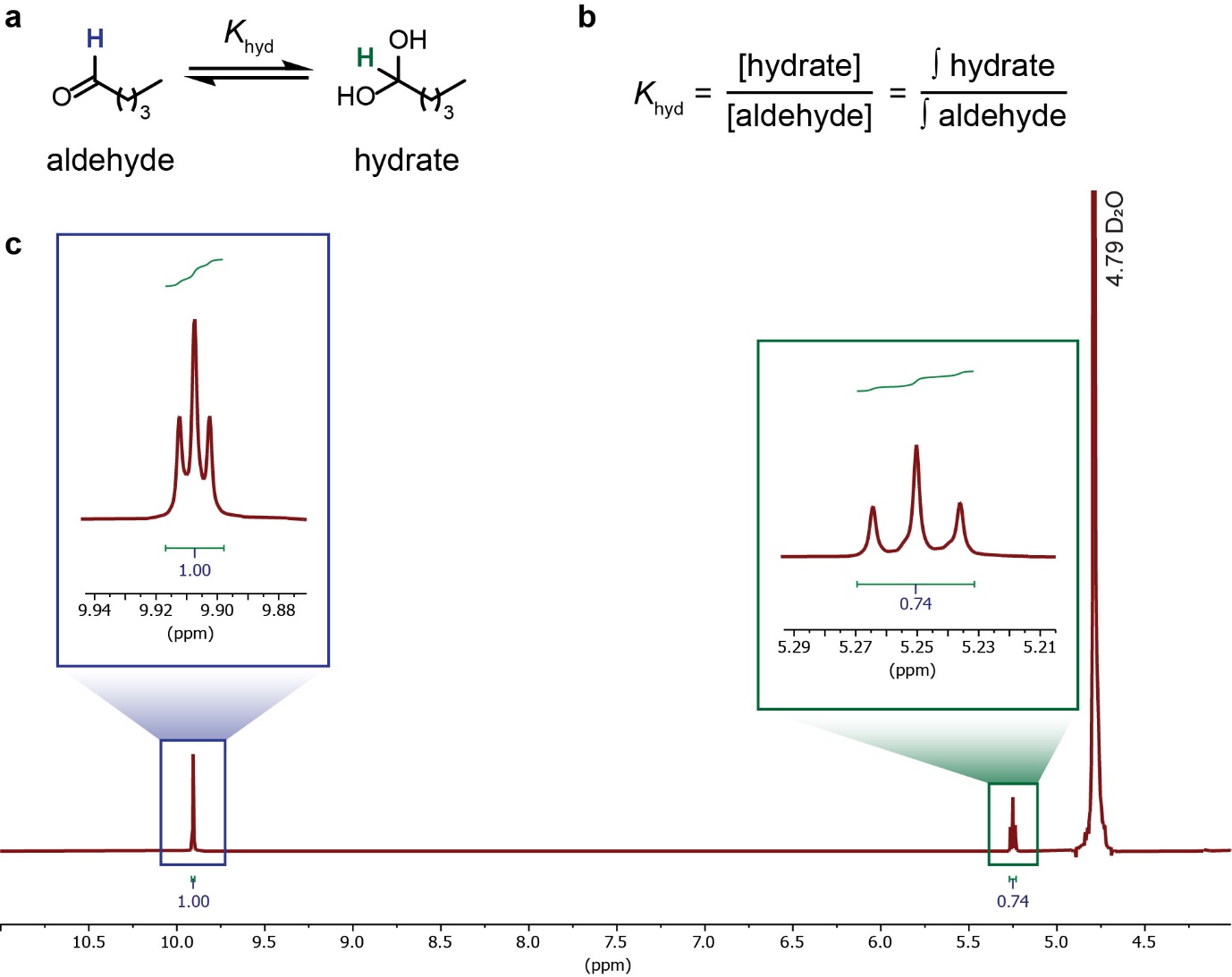

**Supplementary Fig. 11: ^1^H NMR of a pentanal solution.** a) Pentanal is hydrated in aqueous solution, reducing the concentration of free aldehyde available for hemithioacetal formation. b) The hydration equilibrium constant *K*_hyd_ is defined as the concentration of hydrate over the concentration of aldehyde. *K*_hyd_ can be calculated from the ratio of the integration of the peaks for the hydrate gem-diol proton (∫_hydrate_, gem-diol proton colored green) and that of the peaks for the aldehyde proton (∫_aldehyde_, aldehydic proton colored blue). c) ^1^H NMR of 25 mM pentanal in a D_2_O solution containing 2 M KCl and 200 mM PIPES, adjusted to pH 6.4. Integrations of the peaks for the aldehyde proton (boxed in blue) and the peaks for the gem-diol proton (boxed in green) were used to calculate *K*_hyd._

| **Supplementary Table 2:** Hydration constants of aldehyde analytes derived from NMR studies. | | | |
| --- | --- | --- | --- |
| **Aldehyde** | **∫_aldehyde_** | **∫_hydrate_** | ***K*_hyd_ ^[a]^** |
| Ethanal | 1.00 | 1.30 | 1.30 |
| Propanal | 1.00 | 1.18 | 1.18 |
| Butanal | 1.00 | 0.87 | 0.87 |
| Pentanal | 1.00 | 0.74 | 0.74 |
| Hexanal | 1.00 | 0.72 | 0.72 |
| Heptanal | 1.00 | 0.61 | 0.61 |
| 2-Methylpropanal | 1.00 | 0.79 | 0.79 |
| Benzaldehyde | 1.00 | 0 | 0.00 |
| Phenylacetaldehyde | 1.00 | 3.02 | 3.02 |
| ^[a]^ *K*_hyd_ was calculated using ∫_hydrate_/∫_aldehyde_. | | | |

| **Supplementary Table 3:** Concentrations of unhydrated aldehyde in saturated solutions determined by ^1^H NMR. | | | | |
| --- | --- | --- | --- | --- |
| **Aldehyde** | **∫_aldehyde_** | **∫_calcium formate_** | **∫_maleic acid_** | **Saturation concentration (mM) ^[a]^** |
| Hexanal | 1.00 | 0.52 | 0.53 | 19 |
| Heptanal | 1.00 | 2.23 | 2.44 | 4.3 |
| Octanal | 1.00 | 6.74 | 8.74 | 1.3 |
| ^[a]^ Saturation concentrations of each aldehyde were calculated by:  0.5 × (2 × ∫_aldehyde_/∫_calcium formate_ + 2 × ∫_aldehyde_/∫_maleic acid_) × 5 mM | | | | |

### **5 Machine learning**

Events were detected using the single-channel-search function in Clampfit 10.7. Using the start and end times for each event, the event features were extracted using an in-house python script. The dataset was curated from individual events from separate traces of each aldehyde. A total of 9732 events were collected with >900 events per aldehyde (Supplementary Fig. 12).

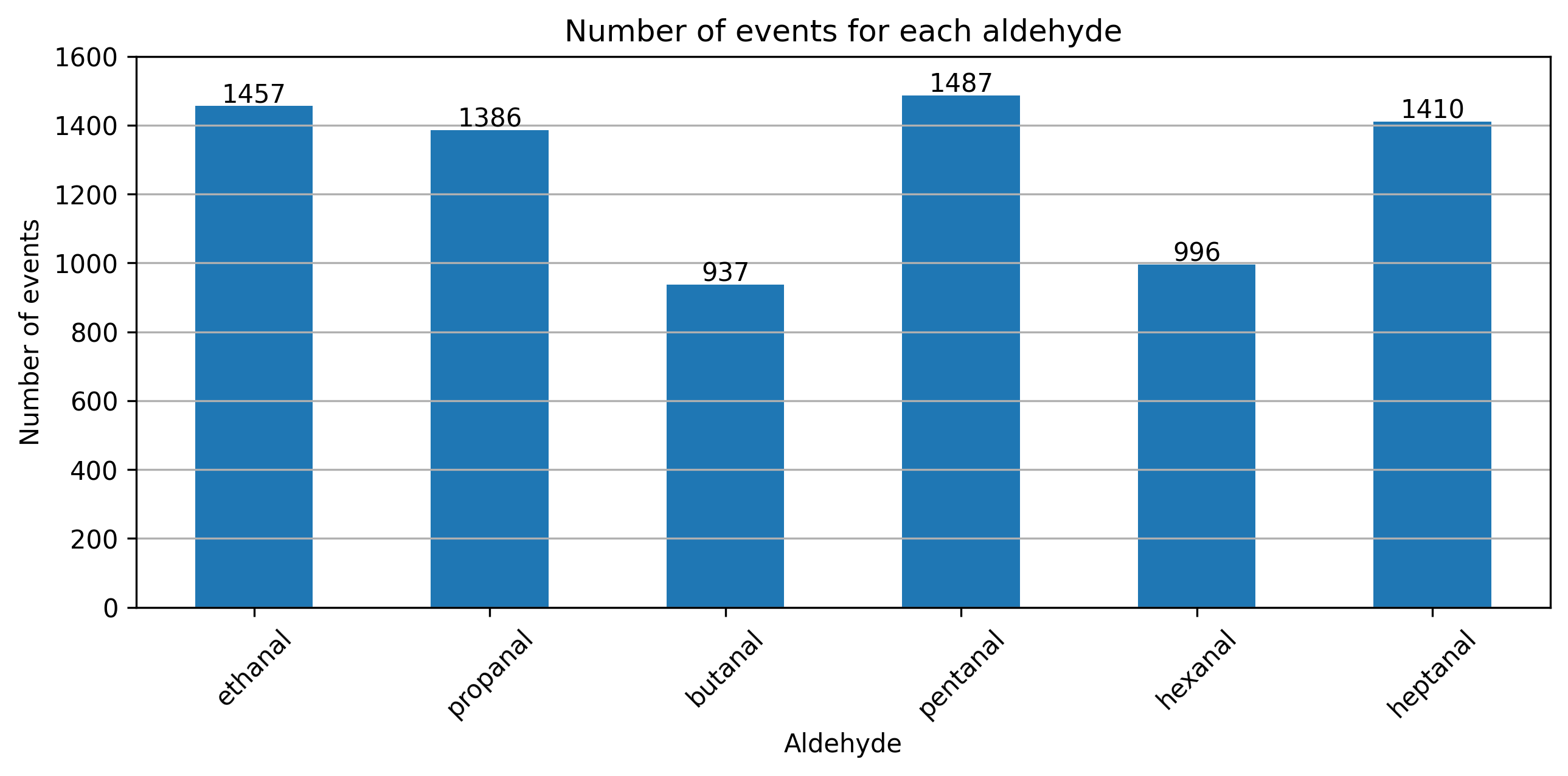

**Supplementary Fig. 12: Events collected for each aldehyde for dataset.**

Three features were extracted for each event, namely the percentage residual current (I_res%_), dwell time, and noise. I_res%_ of each event was calculated as the event’s residual current as a percentage of the preceding open pore current (I_res%_ = I_res_/I_P_ × 100 %). The event dwell time was calculated as the difference between the start and end times of each event. The event noise was calculated as the root mean squared error of the current amplitudes of each event.

A random split stratified on aldehyde labels was performed; 70% of the events were used as training data and the other 30% of events were saved as a test set to test the generality of the model. Five classifiers were tested, including k-nearest neighbors (KNN), decision tree, support vector machine (SVM), random forest, and extra trees (Supplementary Fig. 13). Accuracies were calculated as the ratio of correctly classified event over all events in the dataset. Based on 10-fold cross validation, the random forest model achieved the highest accuracy of 98%. We tested this model on the test set, which also yielded an accuracy of 98%. Source data and codes needed to reproduce the results can be found at: <https://github.com/allybo/aldehyde_nanopore_sensing>.

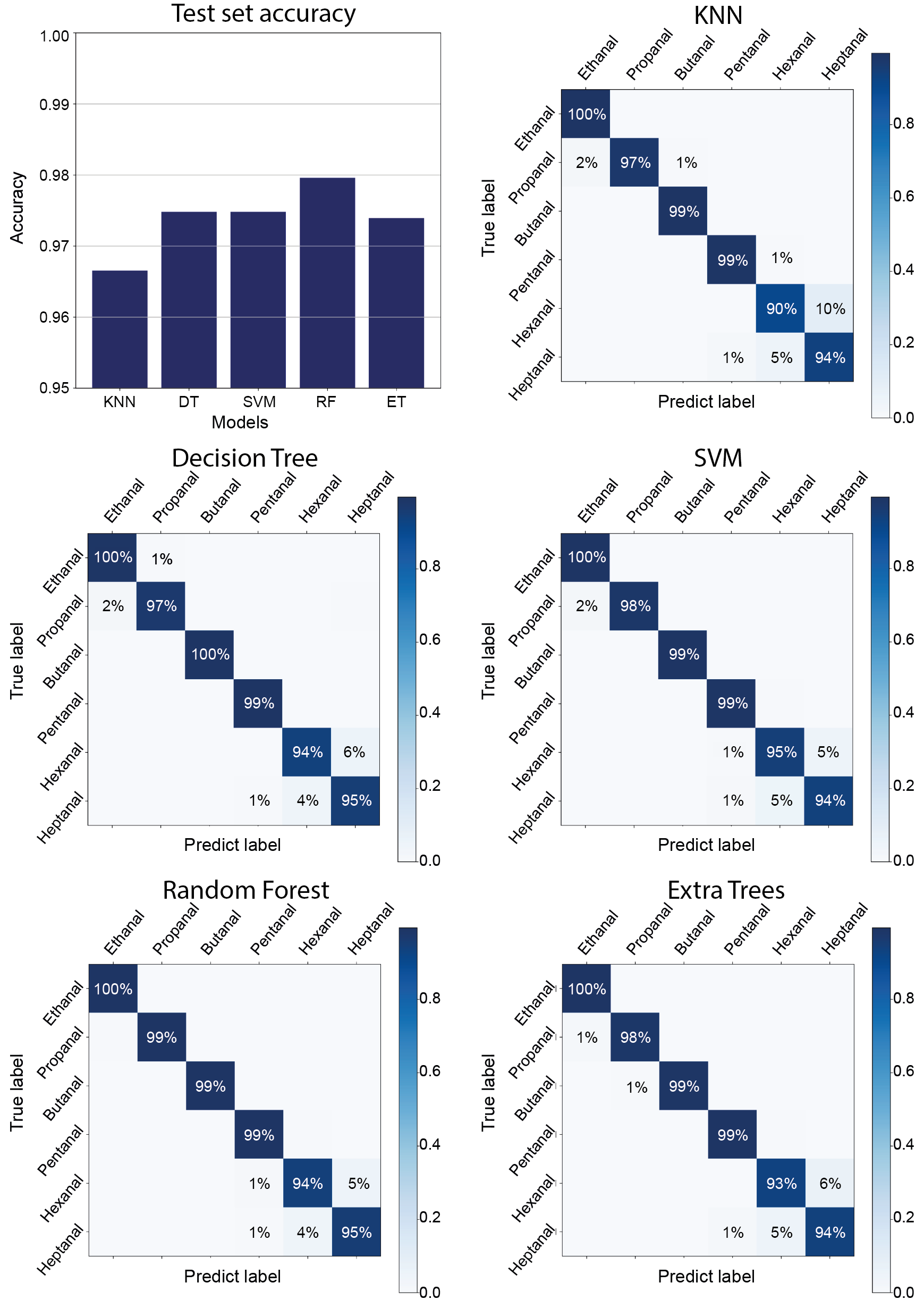

**Supplementary Fig. 13: Model performance of the five classifiers on the test set.** The classifiers tested were k-nearest neighbors (KNN), decision tree, support vector machine (SVM), random forest, and extra trees. Accuracies were calculated as the ratio of correctly classified datapoints over all datapoints in the dataset.

### **6 Rationale for improved diastereomer discrimination in the (AG)_6_(AG-G137C) and (AG)_6_(AG-G137C-Ala3) nanopores**

Nanopores with a cysteine at the position 137 produced larger differences in I_res%_ between structural isomers and diastereomers (compared to nanopores with a cysteine at the 115 position). We performed molecular dynamics (MD) simulations of the (AG)_6_(AG-G137C) and (AG)_6_(AG-G137C-Ala3) nanopores to explore the improved diastereomer discrimination in these nanopores.

MD simulations were performed with GROMACS (v2019.2).^3^ For each system, the nanopore was modeled with the AMBER99SB-ILDN force field and inserted into a cubic box of TIP3P water molecules.^4,5^ The system was neutralized and 0.15 M NaCl was added. Energy minimization was performed with the steepest descent algorithm until the maximum force was below 1000 kJ mol^-1^ nm^-1^. A positional restraint of 1000 kJ mol^-1^ nm^2^ was applied on the protein backbone atoms. Three independent repeats were performed with random initial velocities generated according to the Maxwell–Boltzmann distribution at 298 K. The system was equilibrated under a constant volume and temperature (NVT) ensemble at 298 K (100 ps, 2 fs timestep). This was followed by an equilibration under a constant pressure and temperature (NPT) ensemble at 1 bar and 298 K (100 ps, 2 fs timestep). Production simulations were performed in the NPT ensemble at 1 bar and 298 K (200 ns, 2 fs timestep). The temperature of the system was maintained at 298 K using the V-rescale thermostat.^6^ The pressure was controlled by the Parrinello-Rahman barostat at 1.0 bar, with an isothermal compressibility of 4.5 × 10^-5^ bar^-1^.^7^ All simulations were performed with three-dimensional periodic boundary conditions. Long-range electrostatics was described with the Particle Mesh Ewald (PME) algorithm.^8,9^ All bond lengths involving hydrogen atoms were constrained using the LINCS algorithm.^10^ All three replicas of 50 ns MD trajectories for a given mutant were concatenated.

We assessed the internal pore radius of each mutant using the HOLE program with the default parameters.^11,12^ The narrowest constriction in the wild-type αHL nanopore is formed from the residues of Glu-111, Met-113 and Lys-147. After mutations at these positions (i.e., M113G and K147G) in the (AG)_6_(AG-G137C) and (AG)_6_(AG-G137C-Ala3) nanopore mutants, we observed the removal of the original constriction. Instead, the narrowest part of the (AG)_6_(AG-G137C) and (AG)_6_(AG-G137C-Ala3) nanopore is formed by a ring of leucine residues at position 135. Improved diastereomer discrimination at the 137 position is hypothesized to arise from the location’s proximity to the narrowest constriction site.

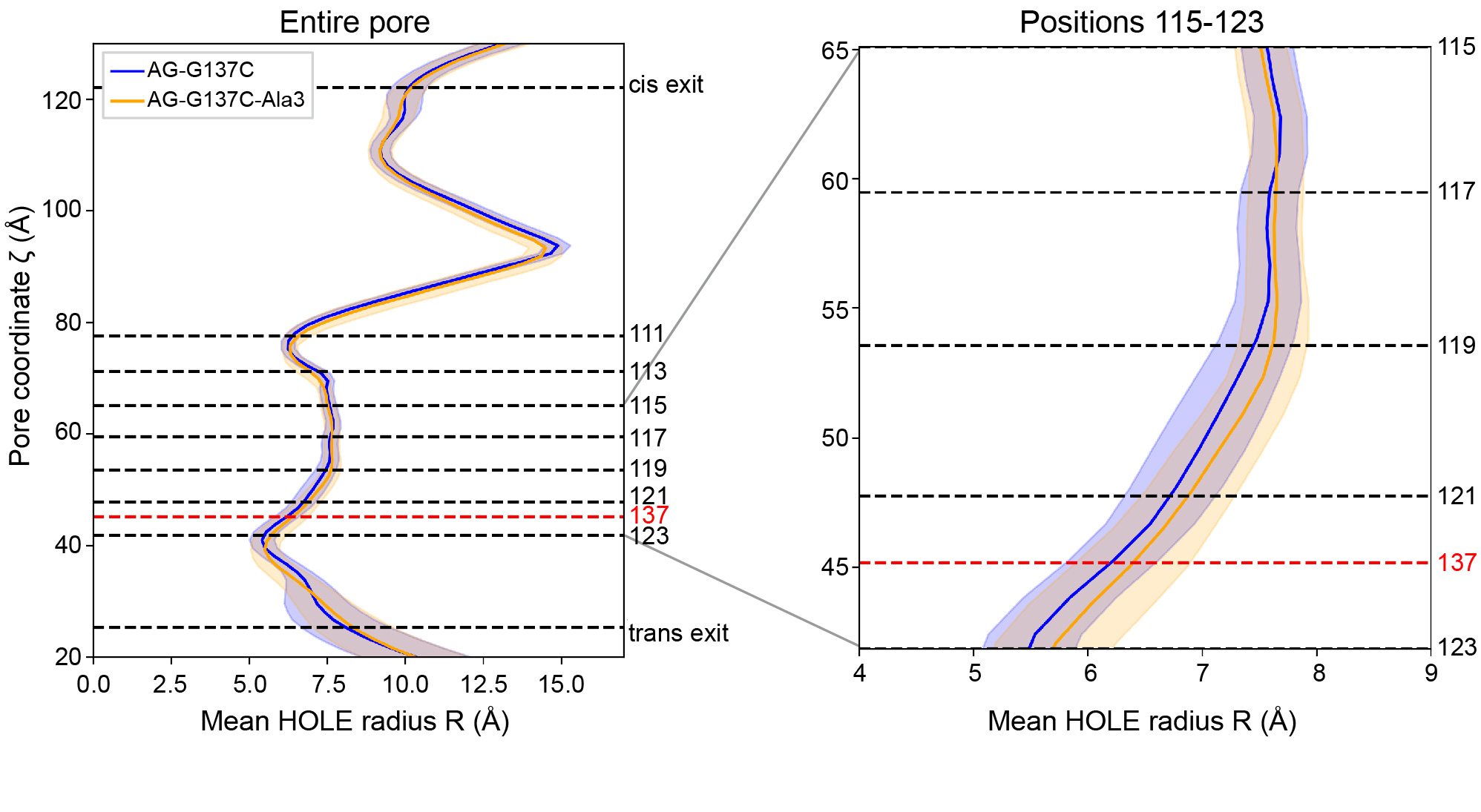

**Supplementary Fig. 14: Model of the internal radius of the nanopore.** Profile of the pore radius along the AG-G137C and AG-G137C-Ala3 nanopores along the z-coordinate. For each mutant, the solid line represents the mean radius, and the shaded region represents the standard deviation.

### **7 Improved structural resolution in engineered nanopore mutants**

Two additional nanopores were designed to address the challenge in distinguishing between structural isomers butanal and 2-methylpropanal. Using the (MK)_6_(MK-T115C) and (AG)_6_(AG-G137C) nanopores, hemithioacetal adducts from ethanal to octanal were easily separable (Fig. 4 and Supplementary Tables 4 & 5).

| **Supplementary Table 4:** Percentage residual current of hemithioacetal adducts of straight-chain aldehydes in (MK)_6_(MK-T115C) nanopore.^[a,b]^ | | |
| --- | --- | --- |
| **Aldehyde** | | **I_res%_** |
| Ethanal | (A) | 99.5 ± 0.1 |
|  | (B) | 99.2 ± 0.1 |
| Propanal | (A) | 98.9 ± 0.1 |
|  | (B) | 98.5 ± 0.1 |
| 2-Methylpropanal | (A) | 98.0 ± 0.1 |
|  | (B) | 97.9 ± 0.1 |
| Butanal | (A) | 97.9 ± 0.1 |
|  | (B) | 97.6 ± 0.1 |
| Pentanal | (A) | 96.9 ± 0.1 |
|  | (B) | 96.5 ± 0.1 |
| Hexanal | (A) | 95.7 ± 0.1 |
|  | (B) | 95.1 ± 0.1 |
| Heptanal | (A) | 94.7 ± 0.1 |
|  | (B) | 93.7 ± 0.1 |
| Octanal | (A) | 93.2 ± 0.1 |
|  | (B) | 92.4 ± 0.1 |
| ^[a]^ Conditions: (MK)_6_(MK-T115C) with 2 M KCl, 20 µM EDTA, 200 mM PIPES (pH 6.8), recorded at −50 mV (trans) and 22 ± 1 °C.  ^[b]^ Errors are standard deviations of the mean values from N = 2 nanopores. | | |

| **Supplementary Table 5:** Percentage residual current of hemithioacetal adducts of straight-chain aldehydes in (AG)_6_(AG-G137C) nanopore.^[a,b]^ | | |
| --- | --- | --- |
| **Aldehyde** | | **I_res%_** |
| Ethanal | (A) | 98.6 ± 0.1 |
|  | (B) | 98.2 ± 0.1 |
| Propanal | (A) | 97.9 ± 0.1 |
|  | (B) | 97.2 ± 0.1 |
| 2-Methylpropanal | (A) | 96.8 ± 0.1 |
|  | (B) | 96.2 ± 0.1 |
| Butanal | (A) | 96.8 ± 0.1 |
|  | (B) | 95.9 ± 0.1 |
| Pentanal | (A) | 95.8 ± 0.1 |
|  | (B) | 94.9 ± 0.1 |
| Hexanal | (A) | 95.9 ± 0.1 |
|  | (B) | 94.1 ± 0.1 |
| Heptanal | (A) | 95.1 ± 0.1 |
|  | (B) | 93.5 ± 0.1 |
| Octanal | (A) | 94.0 ± 0.3 |
|  | (B) | 92.6 ± 0.1 |
| ^[a]^ Conditions: (AG)_6_(AG-G137C) with 2 M KCl, 20 µM EDTA, 200 mM PIPES (pH 6.8), recorded at −50 mV (trans) and 22 ± 1 °C.  ^[b]^ Errors are standard deviations of the mean values from N = 3 nanopores, excluding octanal (N = 2 pores). | | |

Using the (AG)_6_(AG-G137C) nanopore, butanal and 2-methylpropanal can be distinguished using diastereomers B (i.e., ΔI_res%_ of 0.25 ± 0.02 % for diastereomers B of butanal and 2-methylpropanal, N = 3 pores, >35 events for each aldehyde) (Supplementary Fig. 15).

Further engineering was performed on the (AG)_6_(AG-G137C) nanopore to give the (AG)_6_(AG-G137C-Ala3) nanopore (i.e., AG-G137C-Ala3 = WT-K8A-M113G-N121A-N123A-K131G-G137C-N139A-K147G). In the (AG)_6_(AG-G137C-Ala3) nanopore, butanal and 2-methylpropanal could be distinguished using diastereomers A (i.e., ΔI_res%_ = 0.42 ± 0.03 % for diastereomers A of butanal and 2-methylpropanal, N = 4 pores) (Supplementary Fig. 16 and Supplementary Table 6).

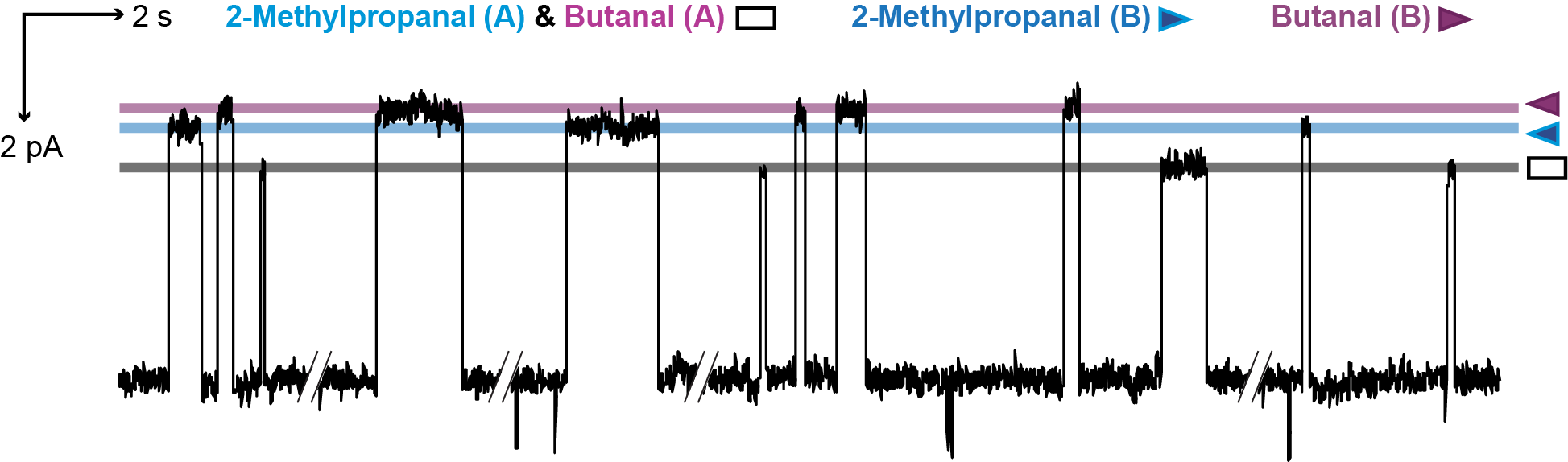

**Supplementary Fig. 15: Structural isomer resolution in the (AG)_6_(AG-G137C) nanopore.** Structural isomers butanal and 2-methylpropanal were distinguished by diastereomers B in the (AG)_6_(AG-G137C) nanopore (i.e., ΔI_res%_ = 0.25 ± 0.02 % for diastereomers B of butanal and 2-methylpropanal, N = 3 pores, >35 events for each aldehyde). Signals were low-pass filtered at 10 kHz and sampled at 50 kHz. Traces were further filtered at 50 Hz for display. Current levels A and B correspond to the diastereomeric adducts.

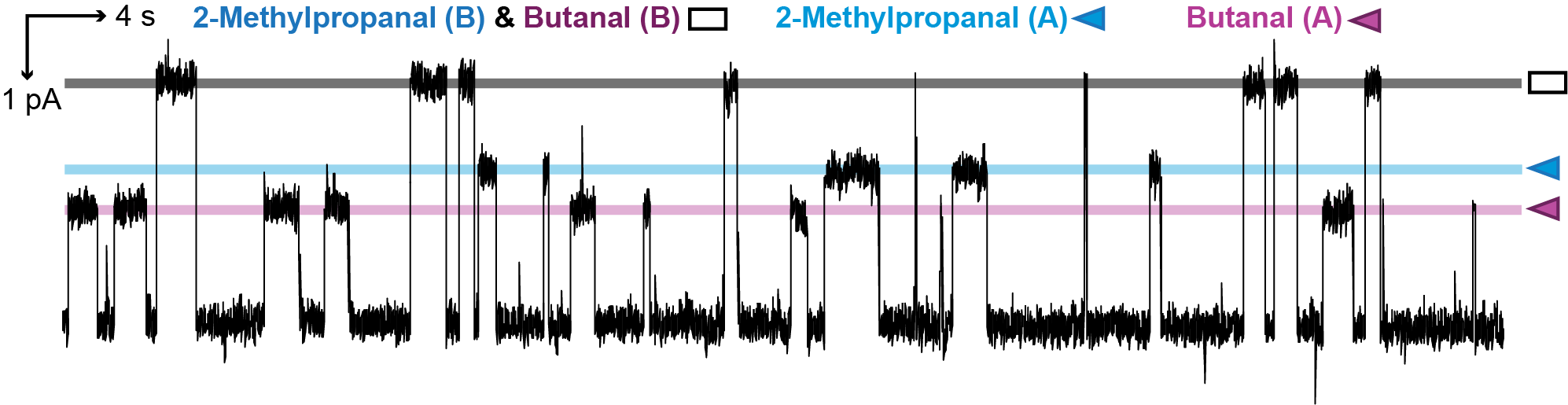
**Supplementary Fig. 16: Structural isomer resolution in the (AG)_6_(AG-G137C-Ala3) nanopore.** Structural isomers butanal and 2-methylpropanal were distinguished by diastereomers A in the (AG)_6_(AG-G137C-Ala3) nanopore (i.e., ΔI_res%_ = 0.42 ± 0.03 % for diastereomers A of butanal and 2-methylpropanal, N = 4 pores, >35 events for each aldehyde). Signals were low-pass filtered at 10 kHz and sampled at 50 kHz. Traces were further filtered at 50 Hz for display. Current levels A and B correspond to the diastereomeric adducts.

| **Supplementary Table 6:** Apparent rate constants of formation (*k*_on_) and dissociation (*k*_off_) of hemithioacetal adducts of structural isomers in (AG)_6_(AG-G137C-Ala3).^[a,b]^ | | | | |
| --- | --- | --- | --- | --- |
| **Aldehyde** | | ***k*_on_ (mM^-1^s^-1^)** | ***k*_off_ (s^-1^)** | **I_res%_** |
| Butanal | (A) | 0.052 ± 0.008 | 0.82 ± 0.07 | 98.6 ± 0.1 |
|  | (B) | 0.030 ± 0.002 | 1.8 ± 0.2 | 97.2 ± 0.2 |
| 2-Methylpropanal | (A) | 0.016 ± 0.002 | 0.86 ± 0.07 | 98.2 ± 0.1 |
|  | (B) | 0.018 ± 0.002 | 2.5 ± 0.4 | 97.1 ± 0.1 |
| ^[a]^ Conditions: (AG)_6_(AG-G137C-Ala3) with 2 M KCl, 20 µM EDTA, 200 mM PIPES (pH 6.8), recorded at −50 mV (trans) and 22 ± 1 °C.  ^[b]^ Errors are standard deviations of the mean values from N = 3 nanopores. | | | | |

### **8 Experimental methods**

#### **General**

All reagents were purchased from Sigma-Aldrich and used without further purification. Catalogue numbers together with purities of aldehydes are listed as follows: Ethanal (≥99.5% purity, Sigma Aldrich catalogue no. 402788); propanal (≥98.0% purity, Sigma Aldrich catalogue no. 64409); butanal (≥99.5% purity, Sigma Aldrich catalogue no. 418102); pentanal (≥98.0% purity, Sigma Aldrich catalogue no. 8.08504); hexanal (≥95.0% purity, Sigma Aldrich catalogue no. 18109); heptanal (≥97.0% purity, Sigma Aldrich catalogue no. 61696); octanal (≥98.0% purity, Sigma Aldrich catalogue no. 52466); 2-methylpropanal (≥99.0% purity, Sigma Aldrich catalogue no. 240788); benzaldehyde (≥99.0% purity, Sigma Aldrich catalogue no. B1334); phenylacetaldehyde (≥95.0% purity, Sigma Aldrich catalogue no. W287407). 1,2-Diphytanoyl-sn-glycerol-3-phosphocholine (DPhPC) used for single-channel electrical recordings was purchased from Avanti Polar Lipids.

#### **Plasmid preparation**

The construction of plasmids encoding the AG (αHL-K8A-M113G-K131G-K147G) and AG-D8 (αHL-K8A-M113G-K131G-K147G-D8) monomers were previously described.^1^

Plasmids encoding the AG-T115C (αHL-K8A-M113G-T115C-K131G-K147G-D8) monomer were prepared from AG-D8 plasmids by a single round of site-directed mutagenesis (Agilent QuikChange II XL Site-Directed Mutagenesis Kit), as follows. Polymerase chain reactions (PCR) (25 µL) were set up by mixing the following reagents in the order listed: 10× reaction buffer (2.5 µL), nuclease-free water (to make up the final volume of the reaction to 25 μL), double-stranded plasmid DNA template (AG-D8 plasmid, 5 µL, 1 ng/µL), mutagenic primers (sense primer: 5′ GAGTATGGGAGTTGTTTAACTTATGGATTCAACGGTAATG 3′; antisense primer: 5′ CATTACCGTTGAATCCATAAGTTAAACAACTCCCATACTC 3′; 62.5 ng of each primer, target mutations are underlined and highlighted in red), dNTP mix (0.5 µL), Quik Solution (1.5 µL) and Pfu Ultra HF DNA polymerase (0.5 µL, 2.5 U/μL). The PCRs were carried out with the following program: 95 °C for 5 min, 18 cycles of: 95 °C (50 s), 53 °C (50 s), 68 °C (5 min), followed by a final extension at 68 °C for 7 min. The PCRs were then cooled on ice for 2 min and the template was digested with DpnI (0.5 μL, supplied with the kit) at 37 °C for 1 h. Plasmids containing the mutant genes were produced from *Escherichia coli* (*E. coli*) XL-10 Gold ultracompetent cells transformed with the PCR product. The DNA sequences of the genes were verified (Source BioScience).

A plasmid encoding the AG-G137C (αHL-K8A-M113G-K131G-G137C-K147G-D8) monomer was prepared from AG-D8 plasmids using a single round of site-directed mutagenesis (Agilent QuikChange II XL Site-Directed Mutagenesis Kit) as follows. A PCR (25 µL) was set up by mixing the following reagents in the order listed: 10× reaction buffer (2.5 µL), nuclease-free water (to make up the final reaction volume to 25 μL), double-stranded plasmid DNA template (AG-D8 plasmid, 5 µL, 1 ng/µL), mutagenic primers (sense primer: 5′ GAATTGGCGGCCTTATTTGTGCAAATGTTTC 3′; antisense primer: 5′ GAAACATTTGCACAAATAAGGCCGCCAATTC 3′; 62.5 ng of each primer, target mutations are underlined and highlighted in red), dNTP mix (0.5 µL), QuikSolution (1.5 µL) and PfuUltra HF DNA polymerase (0.5 µL, 2.5 U/µL). PCR was carried out with the following program: 95 °C for 5 min, 18 cycles of 95 °C (50 s), 60 °C (50 s), 68 °C (5 min), followed by a final extension at 68 °C for 7 min. The PCR was then cooled on ice for 2 min and then the template DNA was digested with DpnI (0.5 μL, supplied with the kit) at 37 °C for 1 h. Plasmids containing the mutant genes were produced from *E. coli* XL-10 Gold ultracompetent cells transformed with the PCR products. DNA sequences of the genes were verified (Source BioScience).

A plasmid encoding the AG-G137C-Ala3 (αHL-K8A-M113G-N121A-N123A-K131G-G137C-N139A-K147G-D8) monomer was prepared from AG-D8 plasmids by four successive rounds of site-directed mutagenesis (Agilent QuikChange II XL Site-Directed Mutagenesis Kit), as follows. Each PCR (25 µL) was set up by mixing the following reagents in the order listed: 10× reaction buffer (2.5 µL), nuclease-free water (to make up the final reaction volume to 25 μL), double-stranded plasmid DNA template (5 µL, 1 ng/µL), mutagenic primers (62.5 ng of each primer, see below), dNTP mix (0.5 µL), QuikSolution (1.5 µL) and PfuUltra HF DNA polymerase (0.5 µL, 2.5 U/μL). PCRs were carried out with the following program: 95 °C for 5 min, 18 cycles of 95 °C (50 s), 60 °C (50 s for all mutants), 68 °C (5 min), followed by a final extension at 68 °C for 7 min. The PCRs were then cooled on ice for 2 min and then the template DNA was digested with DpnI (0.5 μL, supplied with the kit) at 37 °C for 1 h. Plasmids containing the mutant genes were produced from *E. coli* XL-10 Gold ultracompetent cells transformed with the PCR products. DNA sequences of the genes were verified (Source BioScience).

Mutants, primers and DNA templates for each site-directed mutagenesis reaction are shown below. Codons underlined and highlighted in red represent the target mutations in each site-directed mutagenesis PCR.

- **αHL-K8A-M113G-K131G-G137C-K147G-D8** (DNA template: AG-D8; sense primer: 5′ GAATTGGCGGCCTTATTTGTGCAAATGTTTC 3′; antisense primer: 5′ GAAACATTTGCACAAATAAGGCCGCCAATTC 3′)
- **αHL-K8A-M113G-N121A-K131G-G137C-K147G-D8** (DNA template: αHL-K8A-M113G-K131G-G137C-K147G-D8; sense primer: 5′ CTTATGGATTCGCCGGTAATGTTACTGGTGATGATACAGG 3′; antisense primer: 5′ CCTGTATCATCACCAGTAACATTACCGGCGAATCCATAAG 3′),
- **αHL-K8A-M113G-N121A-K131G-G137C-N139A-K147G-D8** (DNA template: αHL-K8A-M113G-N121A-K131G-G137C-K147G-D8; sense primer: 5′ GCGGCCTTATTTGTGCAGCTGTTTCGATTGGTCATAC 3′; antisense primer: 5′ GTATGACCAATCGAAACAGCTGCACAAATAAGGCCGC 3′), and
- **AG-G137C-Ala3** (DNA template: αHL-K8A-M113G-N121A-K131G-G137C-N139A-K147G-D8; sense primer: 5′ GGATTCGCCGGTGCTGTTACTGGTGATGATACAGG 3′; antisense primer: 5′ CCTGTATCATCACCAGTAACAGCACCGGCGAATCC 3′)

Plasmids encoding the MK monomer were prepared from AG plasmids using two parallel rounds of site-directed mutagenesis (Phusion), followed by homologous recombination. The first PCR (25 µL) was set up by mixing the following reagents in the order listed: DMSO (0.75 μL), double-stranded plasmid DNA template (AG, 1 µL, 1 ng/µL), sense primer 5′ GTTTAACTTTAAGAAGGAGATATACATATGGCAGATTCTGATATTAATATTGC 3′ (1.25 μL, 10 μM) and antisense primer 5′ CCGTTGAATCCGTACGTTAACGTACTCATATACTCTTTTGTATCAATCGAATTCCG 3′ (1.25 μL, 10 μM) (target mutations in primers are underlined and highlighted in red), nuclease-free water (8.25 μL), 2 × Phusion Master Mix (12.5 µL). The PCR was carried out with the following program: 98 °C for 30 s, 25 cycles of: 98 °C (10 s), 45 °C (30 s), 72 °C (30 s), followed by a final extension at 72 °C for 7 min. The second PCR (25 µL) was set up by mixing the following reagents in the order listed: DMSO (0.75 μL), double-stranded plasmid DNA template (AG, 1 µL, 1 ng/µL), sense primer 5′ CGGAATTCGATTGATACAAAAGAGTATATGAGTACGTTAACGTACGGATTCAACGG 3′ (1.25 μL, 10 μM) and antisense primer 5′ GCCATATGTATATCTCCTTCTTAAAGTTAAACAAAATTATTTC 3′ (1.25 μL, 10 μM) (target mutations in primers are underlined and highlighted in red), nuclease-free water (8.25 μL), 2 × Phusion Master Mix (12.5 µL). The PCR was carried out with the following program: 98 °C for 30 s, 25 cycles of: 98 °C (10 s), 45 °C (30 s), 72 °C (3 min), followed by a final extension at 72 °C for 7 min. PCR mixtures were then cooled on ice for 2 min and then the template DNA was digested with DpnI (1 μL, 20 U/μL, NEB Catalog #R0176S) at 37 °C for 1 h. Plasmids containing the mutant genes were produced by transforming *E. coli* XL-10 Gold ultracompetent cells (45 μL) with 2 μL of PCR product from each PCR. The DNA sequences of the genes were verified (Source BioScience).

Plasmids encoding the MK-D8 monomer were prepared from AG-D8 plasmids using two rounds of site-directed mutagenesis (Phusion), followed by homologous recombination. The first PCR (25 µL) was set up by mixing the following reagents in the order listed: DMSO (0.75 μL), double-stranded plasmid DNA template (AG-D8, 1 µL, 1 ng/µL), sense primer 5′ GTTTAACTTTAAGAAGGAGATATACATATGGCAGATTCTGATATTAATATTGC 3′ (1.25 μL, 10 μM) and antisense primer 5′ CCGTTGAATCCGTACGTTAACGTACTCATATACTCTTTTGTATCAATCGAATTCCG 3′ (1.25 μL, 10 μM) (target mutations in primers are underlined and highlighted in red), nuclease-free water (8.25 μL), 2 × Phusion Master Mix (12.5 µL). The PCR was carried out with the following program: 98 °C for 30 s, 25 cycles of: 98 °C (10 s), 45 °C (30 s), 72 °C (30 s), followed by a final extension at 72 °C for 7 min. The second PCR (25 µL) was set up by mixing the following reagents in the order listed: DMSO (0.75 μL), double-stranded plasmid DNA template (AG-D8, 1 µL, 1 ng/µL), sense primer 5′ CGGAATTCGATTGATACAAAAGAGTATATGAGTACGTTAACGTACGGATTCAACGG 3′ (1.25 μL, 10 μM) and antisense primer 5′ GCCATATGTATATCTCCTTCTTAAAGTTAAACAAAATTATTTC 3′ (1.25 μL, 10 μM) (target mutations in primers are underlined and highlighted in red), nuclease-free water (8.25 μL), 2 × Phusion Master Mix (12.5 µL). The PCR was carried out with the following program: 98 °C for 30 s, 25 cycles of: 98 °C (10 s), 45 °C (30 s), 72 °C (3 min), followed by a final extension at 72 °C for 7 min. PCR mixtures were then cooled on ice for 2 min and then the template DNA was digested with DpnI (1 μL, 20 U/μL, NEB Catalog #R0176S) at 37 °C for 1 h. Plasmids containing the mutant genes were produced by transforming *E. coli* XL-10 Gold ultracompetent cells (45 μL) with 2 μL of PCR product from each PCR. The DNA sequences of the genes were verified (Source BioScience).

A plasmid encoding the MK-T115C (αHL-K8A-T115C-K131G-K147G-D8; *M*inus-*K*) monomer were prepared from MK-D8 plasmids via a single round of site-directed mutagenesis (Agilent QuikChange II XL Site-Directed Mutagenesis Kit), as follows. Polymerase chain reactions (PCR) (25 µL) were set up by mixing the following reagents in the order listed: 10× reaction buffer (2.5 µL), nuclease-free water (to make up the final volume of the reaction to 25 μL), double-stranded plasmid DNA template (MK-D8, 5 µL, 1 ng/µL), mutagenic primers (sense primer: 5′ CCAAGAAATTCGATTGATACAAAAGAGTATATGAGTTGTTTAACTTATGGATTCAACGG 3′; antisense primer: 5′ CCGTTGAATCCATAAGTTAAACAACTCATATACTCTTTTGTATCAATCGAATTTCTTGG 3′; 62.5 ng of each primer, target mutations are underlined and highlighted in red), dNTP mix (0.5 µL), Quik Solution (1.5 µL) and Pfu Ultra HF DNA polymerase (0.5 µL, 2.5 U/μL). The PCRs were carried out with the following program: 95 °C for 1 min, 18 cycles of: 95 °C (50 s), 55 °C (50 s), 68 °C (5 min 30 s), followed by a final extension at 68 °C for 7 min. The PCRs were then cooled on ice for 2 min and the template was digested with DpnI (0.5 μL, supplied with the kit) at 37 °C for 1 h. Plasmids containing the mutant genes were produced from *Escherichia coli* (*E. coli*) XL-10 Gold ultracompetent cells transformed with the PCR product. The DNA sequences of the genes were verified (Source BioScience).

#### **Nanopore preparation**

(AG)_6_(AG-T115C), (MK)_6_(MK-T115C), (AG)_6_(AG-G137C) and (AG)_6_(AG-G137C-Ala3) heteroheptamers were prepared according to the procedure below, which is a modification of a method previously reported.^13^

αHL monomers were prepared with an *E. coli* *in vitro* transcription and translation (IVTT) system (*E. coli* T7 S30 Extract System for Circular DNA, Cat #L1130, Promega). Prior to use, the T7 S30 extract provided in the kit was treated with 1 µL rifampicin (1 µg/mL, final concentration) to suppress transcription by *E. coli* RNA polymerase. A standard reaction comprised: DNA plasmid mixture (<4 µg, plasmids encoding cysteine-free and cysteine-containing subunits were in an 8:1 ratio), amino acid mixture without methionine (5 µL, as supplied in the kit), S30 premix without amino acids (20 µL, as supplied in the kit), [^35^S]methionine (2 µL, 1200 Ci/mmol, 15 mCi/mL, MP Biomedicals), rabbit red blood cell membranes (2 µL, ~ 1 mg protein/mL), and T7 S30 extract for circular DNA (15 µL, as supplied in the kit). The reaction mixture was incubated at 37°C for 2 h.

Next, MBSA buffer (1 mL; 3-morpholinopropane-1-sulfonic acid (10 mM), NaCl (150 mM), bovine serum albumin (1 mg/mL), pH 7.4) was added to the reaction mixture. The mixture was centrifuged at 12000 RCF for 10 min at 4 °C. The supernatant was then removed, and the pellet solubilized at room temperature with 2× Laemmli sample buffer (25 µL) which contains 10% 2-mercaptoethanol. Before loading the samples, a 5.5 % SDS/PAGE gel was pre-run with 1× Tris-Glycine SDS running buffer containing dithiothreitol (2 mM) and sodium thioglycolate (1 mM). The resuspended pellets were then loaded onto the gel and electrophoresed at +70 V overnight (13 h).

αHL heptamers containing different numbers of mutant subunits were separated in the gel based on their electrophoretic mobilities which were determined by the number of octa-aspartate (D8) tails present (i.e., each cysteine-bearing mutant subunit contained a D8 tail). Hence, the top band corresponded to homoheptamers bearing no octa-aspartate tail (i.e., (AG-αHL)_7_ or (MK-αHL)_7_), the second band corresponded to heteroheptamers bearing a single octa-aspartate tail (i.e., (mutant-D8)_1_(AG-αHL)_6_ or (mutant-D8)_1_(MK-αHL)_6_) and so on, with consecutive bands having an aspartate-tail-free subunit replaced with a mutant subunit bearing an octa-aspartate tail. In this work, the second band from the top contained the desired protein pore with a single cysteine residue.

To extract the protein pores, the gel was dried under vacuum onto Whatman 3MM filter paper for 5 h at 50 °C. The dried gel was then exposed to photographic film (Kodak Bio Max MR autoradiography film) for 8 h and the developed film was used to locate the target protein bands in the gel. The desired protein bands were excised and rehydrated in TE buffer (300 μL; Tris·HCl (10 mM), DTT (0.5 mM), EDTA (1 mM), pH 8.0) for 1 h at room temperature. The paper was then removed, and the gel crushed with a plastic pestle. The resulting suspension was filtered through a 0.2 µm hydrophilic membrane filter (Proteus Mini Clarification Spin Column, Generon). The filtrate was stored in 10 µL aliquots at -80 °C.

#### ***Arthrobacter cholorphenolicus* choline oxidase (AcCO6) preparation**

The plasmid pET28a-AcCO6 encoding an engineered *Arthrobacter cholorphenoculicus* choline oxidase (AcCO6) was a kind gift from Nicholas J. Turner.^17^ The plasmid was transformed in *E. coli* BL21(DE3) competent cells for expression, which were cultured in 10 mL LB medium containing 50 μg/mL kanamycin at 37 ^o^C, 220 rpm overnight. The overnight culture was transferred to 1 L autoinduction media (LB based, Formedium, #AIMLB0110) containing 50 μg/mL kanamycin and grown at 220 rpm for 16 hours at 20 ^o^C. Cells were harvested by centrifugation at 5000 rpm for 20 min at 4 ^o^C. The pellet was resuspended in 50 mL pre-chilled Buffer A (100 mM KPi pH 7.8, 300 mM KCl, 20 mM imidazole) supplemented with 250 U nucleases (Pierce™ Universal Nuclease for Cell Lysis, #88700). Cells were lysed by sonication (10 s pulse, 10 s pause, 3 min duration) on ice, followed by centrifugation at 40000 g for 30 min at 4 ^o^C. The supernatant was filtered through a 0.45 μm syringe filter and applied for purification using a 5 mL HisTrap^TM^ FF column (Buffer A: 100 mM KPi pH 7.8, 300 mM KCl, 20 mM imidazole; Buffer B: 100 mM KPi pH 7.8, 300 mM KCl, 1,000 mM imidazole). Fractions containing His-tagged AcCO6 were pooled and dialysed against a buffer containing 100 mM KPi pH 7.8 at 4 ^o^C overnight. The His-tagged AcCO6 were concentrated, aliquoted, flash frozen with liquid N_2_, and stored at -80 ^o^C.

#### **Single-channel electrical recording**

Single-channel recordings were carried out in a planar bilayer apparatus as previously described.^14^ A single αHL pore was allowed to insert into the bilayer. Aldehyde substrates were introduced from the trans compartment. Experiments were conducted using recording buffer containing 2 M KCl, 200 mM PIPES and 20 μM EDTA titrated to pH 6.8. Aldehyde solutions were prepared with recording buffer and titrated to pH 6.8. Single-channel recordings were conducted with a coverslip placed atop the recording chamber.

Ionic currents were recorded by using a patch clamp amplifier (Axopatch 200B, Axon Instruments), and filtered with a low-pass Bessel filter (80 dB/decade) with a corner frequency of 10 kHz. Signals were digitized with a Digidata 1320A digitizer (Molecular Devices) at an acquisition frequency of 50 kHz. The current traces were processed with Clampfit 10.7 (Molecular Devices). Current traces were idealized by using Clampfit 10.7 (Molecular Devices). The idealized data were analyzed with QuB 2.0 software (www.qub.buffalo.edu).^15^ Dwell time analysis and rate constant determinations were performed by using the maximum interval likelihood (MIL) algorithm of QuB.^16^

#### **Differential sensing of aldehydes and alcohols**

The control aldehyde-alcohol mixture was prepared as follows: propanal, butanal, 1-pentanol, 1-hexanol, 1-heptanol (4 mM each) was dissolved in oxygen-saturated KPi buffer (100 mM, pH 7.8). The treated mixture was prepared as follows: To the control aldehyde-alcohol mixture was added AcCO6 (0.2 mg/mM of alcohol) and allowed to shake overnight at 300 RPM at 30 ^o^C. After incubation, the control or treated mixture was adjusted to pH 6.8, and diluted 4-fold with buffer (2.66 M KCl, 200 mM PIPES at pH 6.8, 20 μM EDTA) (i.e., final concentration of 2 M KCl).

For the enzyme-nanopore coupled experiments, control mixture (100 μL) was first introduced into the trans chamber. As the control mixture was not treated with AcCO6, only propanal and butanal could be detected. Subsequently, treated mixture (100 μL) was introduced into the trans chamber. Addition of the treated mixture produced additional events with conductance levels corresponding to hemithioacetal adducts derived from pentanal, hexanal and heptanal. This confirmed the oxidation of alcohols to aldehydes after AcCO6 treatment.
